# A multiscale theory for mesenchymal cell migration in straight or curved channel confinement

**DOI:** 10.1101/2025.02.26.640464

**Authors:** Wenya Shu, C. Nadir Kaplan

## Abstract

Mesenchymal cells navigate the extracellular matrix (ECM) *in vivo* by processing both its mechanical properties and confinement geometry. Here we develop a multiscale whole-cell theory to investigate cell spreading and migration in two-dimensional (2D) viscoelastic channel confinements of varying width and curvature. Our simulations show that, in straight channels, the cell migration speed depends monotonically on the substrate elastic stiffness, which is otherwise biphasic on an unconfined substrate. This is because confinement enforces directional spreading while reducing the spreading area, which results in lower intracellular viscous drag on the nucleus and a higher net traction force of polarized cells in our model. In contrast, we find that confinement curvature slows down cell migration since the friction forces between the bending cell and the confinement walls increase with curvature. We validate our model with experimental data for cell migration in straight channels spanning a wide range of (ECM) stiffness as well as in curved channels. Our model illuminates the intertwined effects of substrate viscoelasticity and confinement geometry on cell spreading and migration in complex microenvironments, revealing that geometric curvature can hierarchically override substrate mechanics to dominate migration regulation. The study paves the way for designing scaffolds that leverage curvature and confinement to steer controllable cell migration.

**Significance statement:** Cell migration occurs within a 3D extracellular matrix *in vivo*, which confines cell movement into narrow openings. However, most studies focus on migration on open surfaces, overlooking these extra constraints. Here, we develop a theory to reveal how the mechanical properties of the substrate and confinement geometry jointly regulate migration. Our model provides new insights into how cells navigate tissue openings *in vivo* or channels *in vitro*, and highlights how curves along the cells’ trajectories modulate their migration efficiency. Our findings not only improve our understanding of cell motility in development, immune response, and cancer invasion, but also pave the way for the design of biomimetic platforms that control cell movement for tissue engineering and regenerative medicine.

## INTRODUCTION

The extracellular matrix (ECM) is a complex 3D biopolymer network that mediates cell migration *in vivo*. Various experimental and theoretical studies have conventionally regarded the ECM as a 2D tissue culture surface, providing valuable mechanistic insights into how mesenchymal cells form focal adhesions, change shape, and migrate in response to the viscoelastic properties of the ECM [1–6]. However, the 3D ECM topography profoundly modifies the cell behavior from that observed on a 2D substrate, mainly by confining the cells into narrow openings [7–9]. To mimic the effect of 3D confinement on cell mechanics, experiments have commonly used microchannels *in vitro* and demonstrated that channel width, together with ECM elastic stiffness, alters migration behavior [10–14]. Recent experiments also highlighted how channel curvature regulates individual or collective cell migration [15, 16]. To this end, a comprehensive theory that explains the migration dynamics as a function of the channel viscoelasticity, confinement, and curvature can enhance our understanding of mesenchymal cell behavior inside the ECM *in vivo*.

Apart from the channel curvature, the individual effects of the ECM viscoelasticity and confinement on mesencymal migration have theoretically and computationally been investigated [17, 18]. The models rely on the fact that cell migration takes place through sequential, transient lamellipodial or filopodial grips on a substrate by virtue of singular focal adhesions (FAs). The cell gauges the intracellular actomyosin contractile forces to form and reinforce the FAs in the first place, which in turn transmit these forces to the ECM [2, 19, 20]. This dynamic interplay enables cells to actively perceive, react, and adapt to the changes in the ECM biomechanics. The motor-clutch model developed based on this reciprocal relationship between the FAs and actomyosin dynamics successfully elucidated cell adhesion and spreading on substrates with a wide range of elastic moduli and viscosities [4, 21–23]. By spatially connecting several motor-clutch modules to a central cell body, stochastic cell simulators explained the migration behavior on elastic and viscoelastic substrates without actomyosin-regulated FA reinforcement [3, 24–26]. Taking into account confinement, however, necessitates the integration of membrane and cytoskeletal mechanics to rigorously account for the contact forces between the cell and confinement boundaries while maintaining cell structural integrity. Finite element methods (FEM) and phase field models can robustly simulate steric effects associated with confinement barriers, e.g., in the context of adhesion-independent 3D amoeboid migration [27, 28]. Moreover, FEM and cellular Potts models can potentially tackle mesenchymal cell mechanics under confinement as with previous work involving unconfined 2D elastic substrates [29, 30]. Nevertheless, treating the singular profiles of the transient filopodia in FEM as well as modeling viscoelastic media in cellular Potts or phase-field models remain inherently challenging [26, 31, 32]. In contrast, vertex-based or agent-based models that represent the cell by a polygonal mesh provide a straightforward framework to incorporate the contact mechanics between a mesenchymal cell and a rigid barrier while enabling the treatment of substrate viscoelasticity and singular geometries of emergent filopodia during migration [33–36].

Here we generalize our vertex-based multiscale theory in Refs. [35, 36], which were developed for cells migrating on flat, unconfined substrates, to investigate the migration and spreading dynamics in straight or curved channels. Our model integrates subcellular motor-clutch dynamics at the FA sites that form between the cell and the viscoelastic substrate with the structural mechanics of intracellular components and the mechanics of lateral cell contact with channel walls. Our results demonstrate that strong confinement within straight channels significantly suppresses cell spreading but enhances migration speed, in quantitative agreement with experiments [13]. Particularly, on very stiff substrates under strong confinement, the cell spreads less, which alleviates the drag force on the nucleus. This leads to an increase in the net traction force of polarized cells, which in turn promotes the cell migration speed. As a result, the dependence of the migration speed on substrate stiffness becomes monotonic, as opposed to the nonmonotonic profiles observed for unconfined substrates. We also find that increasing channel curvature elevates friction forces between the cell and confinement walls, thereby suppressing both the cell spreading area and migration speed. We further reveal that highly curved channels can even impede cell migration, particularly on viscous substrates with high instantaneous stiffness. Altogether, we integrate the effect of ECM viscoelasticity and confinement geometry into a unified cell migration model that enhances our understanding of cell migration in straight, confined channels and unveils the regulatory role of channel curvature in cell migration.

## METHODS

Cell migration relies on the coordinated interplay of subcellular processes including FA formation, actin polymerization, and actomyosin contraction, as well as the mechanical integrity maintained by structural components such as the cytoskeleton network and cell membrane. In addition to the impact of the ECM mechanical properties on these subcellular mechanisms, geometrical constraints within the microenvironment can directly regulate the deformation and distribution of cell structural components.

Here we extend our multiscale framework [35, 36], originally developed for mechanosensitive cell migration on viscoelastic substrates, to decipher the cell behavior in response to the combined effects of the ECM and confinement width and curvature. Our framework simplifies the migration dynamics to the directional cell spreading accompanied by the nucleus translocation in the absence of inertial effects since cells migrate in low-Reynolds-number environments. At the subcellular scale of our model, the spreading velocity of each vertex, which represents an FA site, is determined by a local force balance between the molecular clutches, membrane forces, and actomyosin contraction (Fig. 1A, B). We employ a chemomechanical coupling between the motor-clutch mechanism and the reaction–diffusion dynamics of the active and inactive Rac1 and RhoA GTPase proteins, whose asymmetric distributions can disrupt the front–rear symmetry of the cell and in turn polarize it. Because of confinement, mechanical equilibrium analysis at a vertex must include a contact detection feature to enforce nonpenetration into the rigid walls (Fig. 1 A), and account for increased contact forces on vertices due to cell bending in highly curved channels (Fig. 1 C). At the cellular level, the traction force at the FA sites, cytoskeletal elastic forces, and contact forces are transmitted to the nucleus. These forces collectively counteract the viscous drag force on the nucleus while the cell moves and drags the nucleus with it (Fig. 1A, D). The speed of nucleus motion is taken as the cell migration speed.

**Figure 1:**
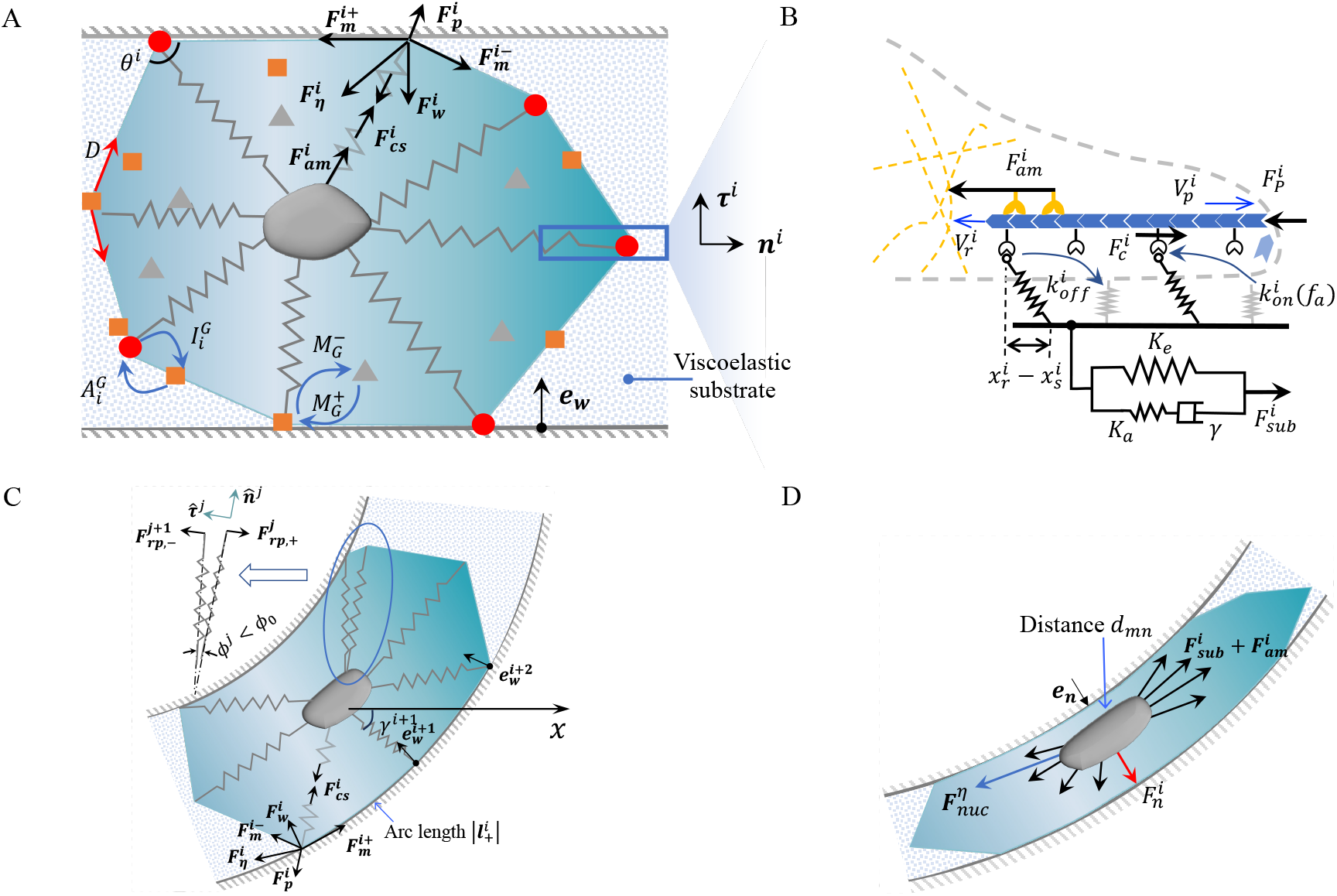
Multiscale model for cell migration on a viscoelastic substrate in confinement. (A) Chemical signaling pathways involve Rho GTPase proteins in active (red dots) or inactive state (orange squares) on the cell membrane, as well as dissolved in the cytoplasm in inactive form (gray triangles). The conversion rates between these states are 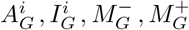 (Eqs S16 and S17, Table S1). *D* : diffusion constant of the membrane-bound species. 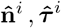: unit radial and polar vectors. (B) Motor-clutch model with stiffness-induced adhesion reinforcement. (C) A migrating cell bent in a curved channel. *ϕ*^*i*^ : angle between two adjacent cytoskeletal springs. When *ϕ*^*i*^ falls below the threshold *ϕ*_0_, repulsive forces 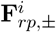 are triggered. 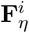 : membrane-wall friction force, 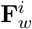 : wall reaction force,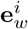: normal vector to the channel wall *γ* angle between the *x*-axis and *i*-th cytoskeletal spring. (D) Forces on the cell nucleus in a highly curved channel. 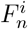 : constraint force acting in the normal direction **e**_*n*_ that prevents the nucleus-membrane distance *d*_*mn*_ from falling below a threshold *d*_*min*_. All other parameters and variables in (B)–(D) are defined in Table 1.

### Equations of motion at subcellular scale

To update a vertex position **x**^*i*^(*t*) in time *t*, the vertex spreading speed 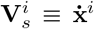 is determined from the local force balance at that vertex when the corresponding FA dynamics is known. The local force balance depends on whether or not a vertex contacts a channel wall. In the following, we first provide an overview of the spreading dynamics at a vertex as it was implemented in our previous models [35, 36]. Then, we detail its generalization that considers dynamic wall contact of a cell under confinement.

**Table 1:**
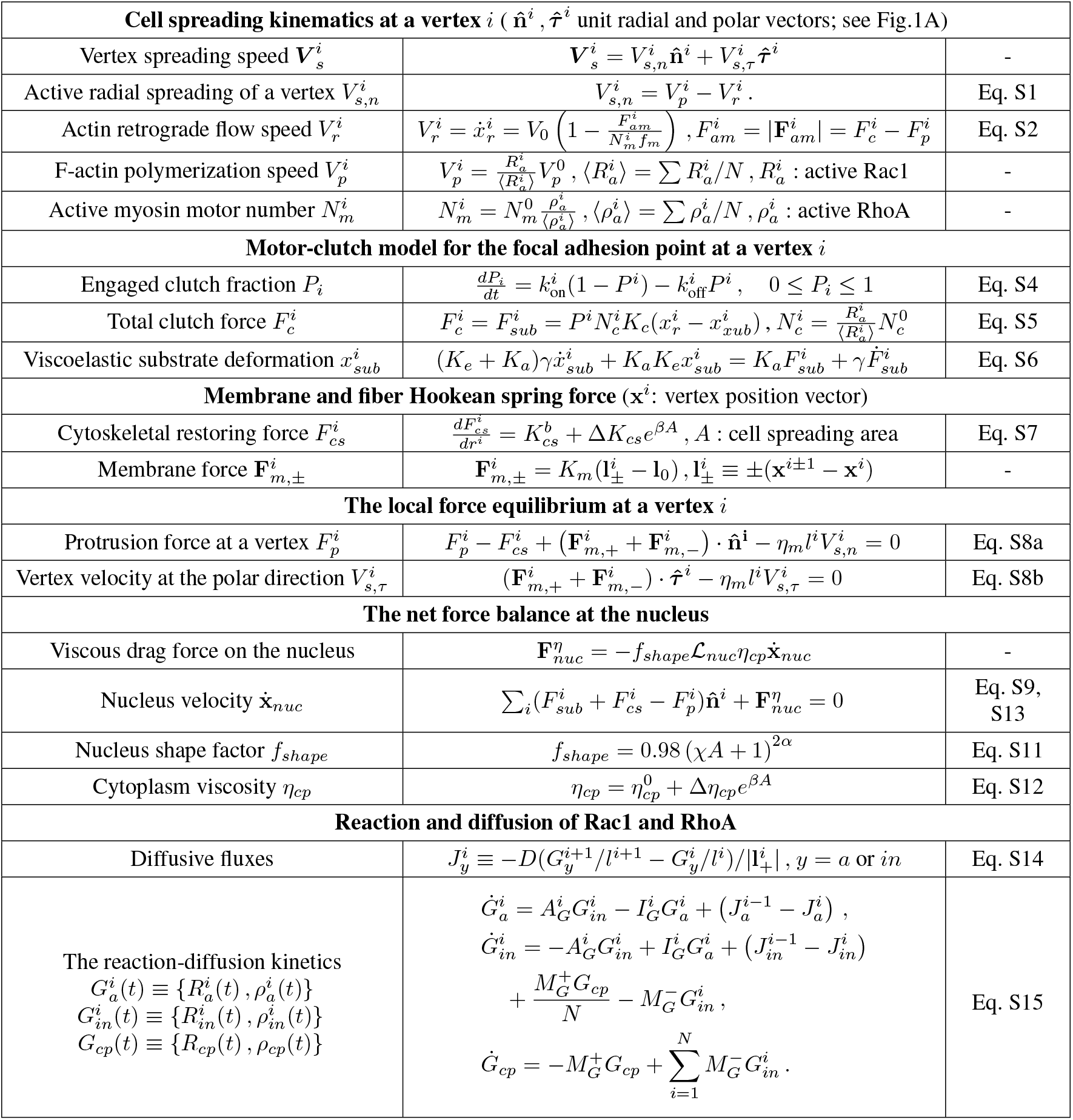
Model summary for cell migration without confinement. *V*_0_ : Unloaded retrograde flow speed, 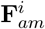 : active pulling force acting on the actomyosin complex, *f*_*m*_ : single myosin motor stall force, 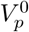: characteristic polymerization rate, *N* : number of vertices, 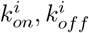 : rate constants of clutch association/dissociation (Sec. S1.1.1), 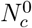 : reference molecular clutch number, 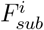 : substrate deformation force, *K*_*c*_ : clutch stiffness, *K*_*e*_, *K*_*a*_, *γ* : SLS model parameters (Fig. 1B, Table 2), 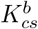 : baseline cytoskeletal stiffness, Δ*K*_*cs*_ : stiffness increment, *β* : Area-dependent nonlinearity coefficient, *K*_*m*_ : membrane stiffness, *α, χ* : shape factor parameters (Sec. S1.2.1), 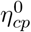: reference cytoplasm viscosity, Δ*η*_*cp*_ : cytoplasm viscosity increment, *D* : membrane-bound rho GTPase diffusion constant, *a* : active, *in* : inactive. For the numerical values of the parameters used in the simulations, see Table S1.

In our model, the radial component of the local spreading velocity 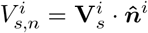 is primarily dictated by the active forces associated with the FA mechanics that is regulated by the Rho GTPases, while the polar component 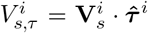 relies on the balance between the polar component of the passive forces due to the cell membrane deformations and steric interactions, such as those from channel walls (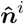and 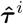 defined in Fig. 1A). The radial spreading speed is set by the competition between the actin polymerization rate 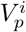 and the retrograde actin flow speed 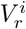, i.e., 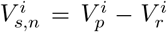 (Fig.1B). We compute 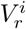 by employing a mean motor-clutch model at each vertex, which considers that proteins such as integrin, talin, and vinculin can form temporary molecular clutches with the ECM to generate adaptive traction while resisting actin retrograde flow (Sec. S1.1 in the Supporting Information) [21–23]. The adhesion reinforcement effect in our motor clutch model relates the clutch association rate 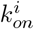 monotonically to the substrate elastic stiffness, which sets the local cell traction force strength 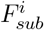 and in turn determines the cell spreading area by affecting 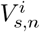 (Sec. S1.1) [36]. To determine the direction of local spreading, we couple 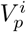 and 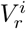 to the Rac1 and RhoA proteins that undergo reaction and diffusion (Sec. S1.2); that way, our model facilitates emergent mechanical polarization of the cell from the dynamical chemical polarity of the Rho GTPases.

To explore how the ECM viscoelastic properties regulate cell spreading and movement, we adopt the standard linear solid (SLS) model for viscoelasticity. The SLS model comprises an elastic spring linked in parallel with a Maxwell arm (see Fig.1B), where *K*_*e*_ represents the residual elastic stiffness as *t* → *∞* after the viscous stresses have relaxed, *γ* is the substrate viscosity, and *K*_*a*_ is the additional stiffness that governs the stress relaxation with a timescale *τ*_*r*_ ≡ *γ/K*_*a*_. In the limit *t* → 0, the SLS model yields an instantaneous stiffness *K*_*t*→0_ = *K*_*e*_ +*K*_*a*_ for the elastic response of the substrate before the viscous relaxation of the stress.

Maintaining the structural integrity of the cell to accurately characterize both the contact forces and substrate-stiffness-dependent cell spreading areas in our simulations requires that we consider force transmission between adjacent vertices and between each vertex and the nucleus (Sec. S1.1). To link the nucleus to the peripheral vertices, our framework represents the cytoskeleton as a set of nonlinear hyperelastic springs that incorporate strain stiffening. To represent each membrane section between adjacent vertices, we consider a linear spring with stiffness *K*_*m*_ and friction constant *η*_*m*_ to account for its friction with the ECM. The vertex local equilibrium condition contains all these components to determine the spreading velocity 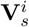 (Table 1). However, when a vertex and the connected cell membrane section contact a rigid channel wall, the steric interactions restrict the vertex movement and concurrently augment the membrane-wall friction force, which significantly modify the local force balance as we describe next.

### Vertex contact with straight channel walls

For a cell migrating in a rigid, straight channel, once the vertex *i* contacts the channel wall, the reaction force from the wall 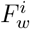 must result in zero vertex speed in the normal direction to the wall 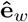 (Fig. 1A). Moreover, contact between the membrane and the wall increases the local friction force. Therefore, we assume an augmented drag coefficient 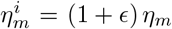 to account for the enhanced friction during the movement of vertex *i*. Additionally, during contact, the membrane itself becomes more resistant to deformations due to a higher friction force. To accommodate this, we introduce an enhanced membrane stiffness denoted as 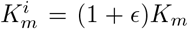. The value *ϵ* = 0.5 was chosen based on experimental fits on the migration speed of U373-MG human glioma cells in straight channels (Sec. S2, Fig. S1) [13]. Then, the following equilibrium equations with the nonpenetration condition at the vertex *i* must hold:

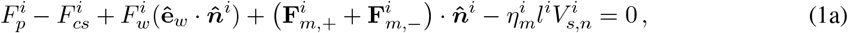

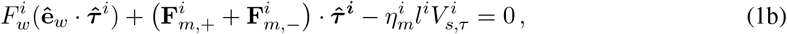

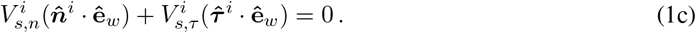

The membrane forces at the *i*^*th*^ vertex are given as 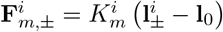. Here, 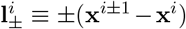 are the distance vectors between vertex *i* and its two neighbors, and **l**_0_ is the initial distance between two adjacent vertices. In Eqs. 1a and 1b, the variable 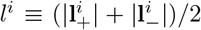 defines the average membrane length about the *i* ^*th*^ vertex. Eq. 1a determines the membrane force resisting protrusion 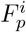 and the radial spreading speed 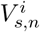 together with Eq. S1, S2, and S5 (Table 1), Eq.1b determines the polar spreading speed 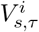, and Eq.1c sets the reaction force from the wall 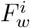 by coupling the active and passive dynamics at vertex *i* to satisfy the nonpenetration condition.

### Vertex contact with curved channel walls

In curved channels, the reaction forces at the cell-wall interface must counterbalance the internal forces that are generated due to the bending of the cells. This is because, under confinement, the mechanicaly stable state of the cell is a straight, elongated configuration along its polarization direction due to its structural integrity, which is collectively maintained by the cytoskeletal network, contractile actomyosin tension, and membrane resistance as included in our model. To test the curvature dependence of these contact forces, we perform FEM simulations of a bending elastic beam that represents an elongated cell within a curved channel (Sec. S3). We find that the reaction forces (equivalent to the contact forces) at the cell-wall interface have a quadratic dependence on the channel curvature as dictated by symmetry under the curvature sign change (Fig. S2). Since our vertex-based model represents the cell as a cable-truss structure without any inherent bending stiffness, we explicitly introduce a curvature-dependent stiffness to the cell membrane as

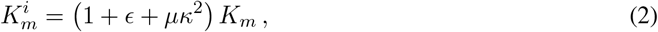

where *µ* = 2 × 10^4^ *µm*^2^ was determined through an iterative model calibration to match the experimental elongation length measurements of MDA-MB-231 cells in a circular channel with a radius *R* = 50 *µm* (see Sec. S2 and Fig. S3) [15]. In straight channels, we introduced an augmented drag coefficient with a fitting parameter *ϵ* above. In curved channels, curvature-dependent reaction forces further amplify the membrane-wall friction during migration. To account for this, we introduce a modified drag coefficient:

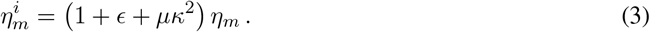

Noting that the modified membrane forces are given by 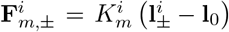 per Eq. 2, the spreading-induced friction forces are given by 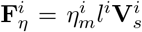 per Eq. 3, and the normal vector of the curved surface changes with the vertex position (e.g., 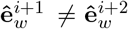 ; Fig. 1C). Then, the force balance at a vertex *i* with a nonpenetration condition in a curved channel is written as

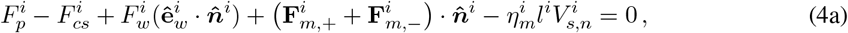

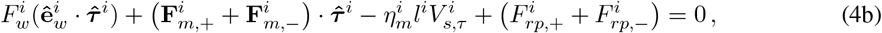

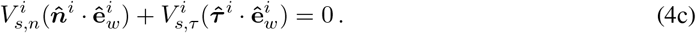

The last term including 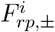 on the left-hand side of Eq. 4b represents repulsive forces in the polar direction. These forces are incorporated to prevent the adjacent cytoskeletal elements from overlapping or crossing in highly curved channels. The computation of these repulsive forces is based on the penalty method from contact mechanics, which implies that the force is applied only when the angle *ϕ*_*i*_ becomes smaller than a threshold value *ϕ*_0_ (Sec. S4). In general, 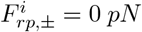 for most vertices.

Eqs. 1a-4c complement Eq. S8 to calculate the spreading velocities 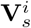 simultaneously for all vertices, which are in or without contact with the channel walls at a given time. That way, the model takes into account the long-range influence of contact forces to distant vertices without contact in that the local forces are transmitted not only to the neighboring vertices through the cell membrane, but also to the distant vertices through the cytoskeletal network over the cell nucleus, as detailed in the following.

### Equation of motion of the nucleus

Our model represents the motion of the cell body by the translation of its nucleus. The linkage between the cytoskeleton and nucleus facilitates the transmission of various forces (Fig. 1A, D), including the cytoskeletal elastic forces 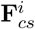 and the active force resisting actomyosin contraction 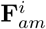 (Fig. 1B, Table 1). The sum of these cytoskeletal and peripheral forces lead to the nucleus deformation and movement [37]. The nucleus velocity 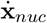 is computed by the balance between these forces and the viscous drag force within the cytoplasm that is given as 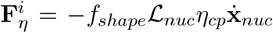 where *f*_*shape*_ is the shape factor that measures the deviation of the nucleus from a perfect sphere with 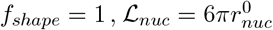 and 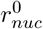 is the radius of the undeformed spherical nucleus, and *η*_*cp*_ is the cytoplasm viscosity (Eq. S9 and Table 1). This relation implies that when the shape factor *f*_*shape*_ = 1, the viscous force reduces to the Stokes drag. Since cell spreading can significantly deform the nucleus and increase the cytoplasm viscosity, we related both the shape factor *f*_*shape*_ and the cytoplasm viscosity *η*_*cp*_ to the spreading area (Eqs. S11-S12, Table 1) [38–40]. This contributes to an augmented viscous drag force on the nucleus with increasing cell spreading area, hampering the cell migration speed on high-stiffness substrates in the absence of confinement [36].

In extremely curved channels, cell bending and elongation in our model may lead to a contact between the nucleus and the cell membrane. This is nonphysical since the nucleus and the cell membrane are typically separated by multiple layers of other cellular components, with the minimum distance between the nucleus surface and the cell surface being typically around 1 *µm* [37, 41]. *Therefore, if the distance between the nucleus and membrane (d*_*mn*_ in Fig.1E) falls below a minimum value of *d*_*min*_, we introduce a constraint force *F*_*n*_ to enforce a zero velocity for the nucleus in the direction normal to the membrane (**e**_*n*_ in Fig. 1E). This gives an updated equilibrium system:

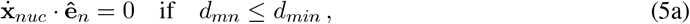

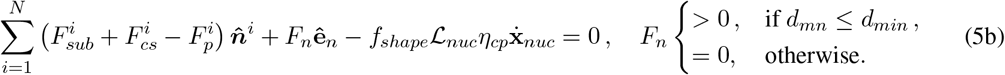

Eq. 5a determines the contact force in Eq. 5b if *d*_*mn*_ ≤ *d*_*min*_, and Eq. 5b yields the nucleus velocity 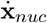.

### Simulation procedure

We represent the initial vertex configuration of the cell as a regular polygon with *N* = 16 sides due to its decent convergence properties as with our previous work [35, 36]. All initial forces, velocities, displacements, and substrate deformations are set to zero. Since chemical signaling drives cell polarization followed by directional locomotion on uniform substrates, we enforce a nonuniform and stochastic initial distribution of the active Rac1 protein volume fraction, 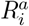, with higher values at random vertices at the cell front to mimic the random nature of the protrusion formations in the mesenchymal cells [35]. Likewise, a polarized and stochastic initial distribution of the active RhoA protein volume fractions 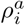 is taken with accumulation at the cell rear (manual polar symmetry breaking). In contrast, the initial conditions for 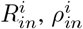 are uniform at time *t* = 0, and the cytoplasmic form *R*_*cp*_, *ρ*_*cp*_ can be determined from the total volume fraction conservation (Sec. S5). This initial condition of Rho GTPase proteins contributes to the polarization and directed cell migration on unconfined substrates and along the direction of the confined channels. Starting from these initial conditions, we update the radial spreading speed of each vertex by solving the mean-field motor-clutch model (Eqs. S1–S6, Table 1). The calculation of the spreading velocity requires a contact condition assessment between a vertex and the channel wall: When a vertex is not in contact with the wall, Eq. S8a,b are used. Otherwise, Eqs. 1a–1c and Eqs. 4a–4c are used for vertex in contact with straight or curved channels, respectively. The nucleus velocity is calculated by solving Eq. S13 or Eq. 5a–5b, depending on its proximity to the cell membrane (*d*_*mn*_ in Fig. 1D). Meanwhile, the Rho GTPase volume fractions are updated by solving the reaction and diffusion system in Eqs. S14, S15 by following the algorithm in Fig. S5. When the cytoplasmic inactive Rac1 volume fraction *R*_*cp*_ reaches a steady state, we reinitialize 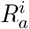 and 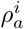 stochastically to mimic the random nature of the protrusion formations in the mesenchymal cells. All fixed simulation parameters are listed in Table S1. The range of channel geometric parameters and ECM viscoelastic properties, which are of interest in this study, are provided in Table 2.

**Table 2:** Material and geometric parameters. The parameters *K*_*e*_, *K*_*a*_, *γ* are defined in Fig. 1B.

| Symbol | Parameters | Dimensional | Dimensionless |
| --- | --- | --- | --- |
| $K_e$ | Elastic stiffness | 0.1-120 pN/nm [13, 21, 22] | $5\pi - 6000\pi$ |
| $K_a$ | Additional stiffness | 1.0 - 20 pN/nm [22] | $50\pi - 1000\pi$ |
| $\gamma$ | Viscosity | 0.1 - 100.0 pN·s/nm [22] | 0.06 - 60 |
| $W$ | Channel width | 8 - 40 $\mu\text{m}$ [13] | 0.4 - 2 |
| $\kappa$ | Channel curvature | 0 - 0.02 1/$\mu\text{m}$ | 0 - 0.4 |

We present the simulation results in dimensionless form by using a position scale *L* ≡ 2*πr*_0_ = 2*π* × 3*µm*, speed scale *V*_0_ = 120 *nm/s* (unloaded retrograde flow speed; Table S1), migration timescale *T* ≡ *L/V* ≈ 160 *s*, and force scale 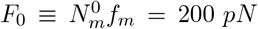. Each simulation is run over *M >* 10^6^ time steps with a time step of Δ*t* ≈ 10^−3^ *s* (Δ*t* ≈ 10^−5^ in dimensionless units), corresponding to the migration dynamics over at least an hour in real units. The spreading area, *A*, and the aspect ratio, *δ*, are determined as the average cell area and the average ratio of the cell’s shortest axis to its longest axis, respectively, calculated over the last 10^6^ time steps of the simulation. We define the migration speed *V* as the nucleus trajectory length *X* ≡|**x**_*nuc*_(*M* Δ*t*) − **x**_*nuc*_(0) | divided by the total time *t*_*total*_ ≡ *M* Δ*t*, i.e., *V* ≡ *L/t*_*total*_. For each set of the ECM parameters and geometric factors (Table 2), we run *n* simulations to compute the sample mean spreading area, 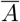, and the mean migration speed, 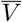. The effects of the ECM parameters and geometric factors on the migration dynamics are validated by the experiments in Refs. [13, 15].

## Results

### Effect of confinement on cell spreading and migration

We first investigate the impact of physical confinement in a straight channel on the areal spreading and migration speed of cells on elastic substrates (i.e., viscosity *γ* → 0). Regardless of channel confinement, cell spreading area on elastic substrates consistently increases with substrate stiffness due to the adhesion reinforcement effect (Fig. 2A). However, since the reaction force 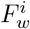 on a vertex that is contact with a channel wall leads to an increasing radial protrusion force 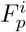 at that vertex (see Eqs. 1a and 4a), the radial spreading speed 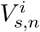 of the vertex decreases (see Eq. S1-S2). As a result, channel confinement significantly suppresses cell spreading, reproducing the experimental data obtained from U373-MG human glioma cells on polyacrylamide hydrogels [13]. This suppression is more pronounced on stiffer substrates because increased substrate stiffness promotes greater cell spreading, resulting in higher reaction forces 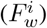 from the channel walls. Consequently, the areal spreading is significantly reduced on stiff substrates 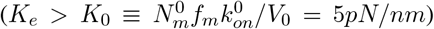 compared to the softer ones (*K*_*e*_ *< K*_0_) (Fig. 2A). Channel confinement not only quantitatively affects migration speed but also qualitatively governs migration behavior (Fig. 2B). Introducing confinement on a stiff substrate boosts the cell migration speed, transforming the biphasic speed-stiffness relationship on unconfined substrates into a monotonic profile. One underlying reason is the suppression of the areal spreading in confined channels, which decreases both the shape factor *f*_*shape*_ (Eq. S11) and the cytoplasm viscosity *η*_*cp*_ (Eq. S12). As a result, the viscous drag coefficient of the nucleus (defined as *f*_*shape*_*η*_*cp*_) is significantly reduced in confined channels, particularly on stiff substrates (Fig. 2C). This reduction in the viscous drag coefficient reduces the viscous drag force on the nucleus, thereby enabling faster migration as substrate stiffness increases in confined channels (Fig. 2D). Additionally, confinement on very stiff substrates (*K*_*e*_ ≫ *K*_0_) not only induces cell elongation within the channel but also aligns the cytoskeletal elements at the front and rear of a migrating cell (see Movie 1). This is supported by the experiments that demonstrate an alignment within the intracellular distribution of phosphorylated myosin light chain and F-actin [13]. As traction forces become increasingly aligned, confinement enhances the net traction force of polarized cells on stiff substrates (Fig. 2D), effectively promoting cell migration speed and contributing to the monotonic speed-stiffness relationship.

**Figure 2:**
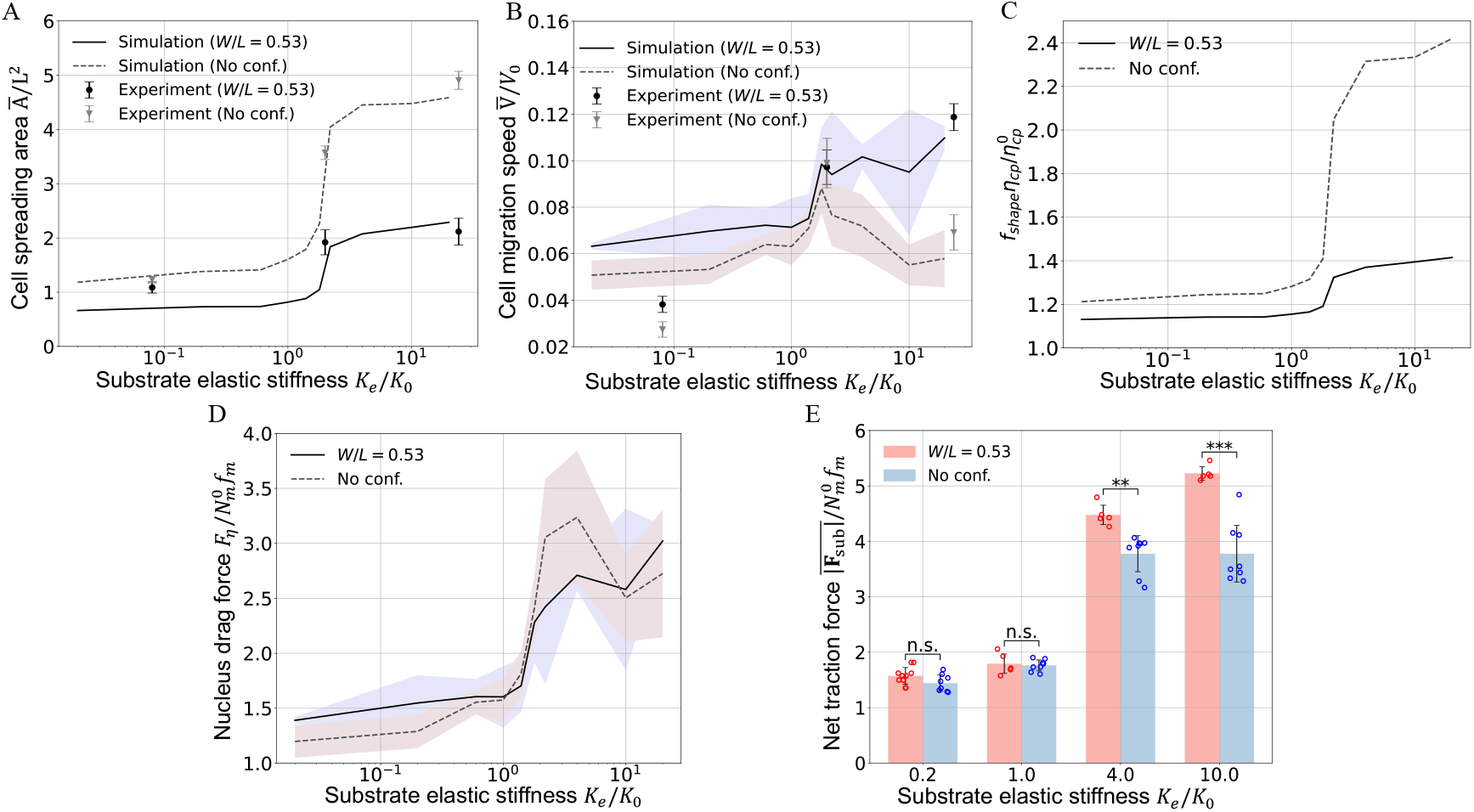
Matrix confinement suppresses cell spreading and promotes migration speed. (A) Unitless mean spreading area *Ā/L*^2^ and its standard deviation versus the uniform unitless elastic stiffness of a substrate, *K*_*e*_*/K*_0_, for cells migrating in a confined channel (full line, *W/L* = 0.53). The characteristic threshold stiffness 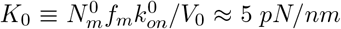 classifies the substrate as ‘stiff’ when *K*_*e*_ *> K*_0_ and ‘soft’ when *K*_*e*_ ≤ *K*_0_ (Sec. S1). The dashed line shows the dependence of *Ā/L*^2^ on the stiffness of an unconfined substrate. (B) Unitless migration speed 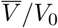 and its standard deviation as a function of the substrate elastic stiffness *K*_*e*_ for fixed channel width *W/L* = 0.53. The dashed line is the biphasic stiffness/mean speed relation on uniform substrates without spatial confinement [36]. (C) Viscous drag coefficient of the nucleus 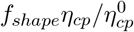 versus *K*_*e*_*/K*_0_ on unconfined substrates and within confined channels. The calculation of *f*_*shape*_ and *η*_*cp*_ depends on the computed spreading area (Eqs. S11, S12, Table 1). (D) Unitless viscous drag force on the nucleus 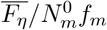 and its standard deviation as a function of the substrate elastic stiffness *K*_*e*_ for fixed channel width *W/L* = 0.53. The dashed line represents the stiffness/nucleus drag force relation on uniform substrates without spatial confinement. (E) Comparison between the mean net traction forces 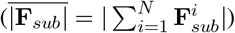 of cells migrating on unconfined substrates and within confined channels as a function of unitless substrate stiffness. Statistical analysis is performed for each pair of results with the statistical significance (*p*) denoted as: ∗∗ ∗: *p <* 0.001, ∗ ∗: *p <* 0.01, ∗: *p <* 0.05, and n.s.: *p >* 0.05 (Sec. S6). In (A) and (B), the mean values and the standard deviations are calculated over *n* = 10 simulations, while dots and error bars show the experimental data from Ref. [13] [42].

We next explore the impact of channel width on cell migration speed at constant substrate stiffness. Note that our simulations employ persistent polarization along the channel wall characterized by a well-defined leading edge with polarized chemical signaling. This is an Ansatz; after all the influence of confinement on chemical polarity remains poorly understood in the existing literature. Our findings demonstrate that narrower channels promote faster migration on stiff and very stiff substrates, in agreement with the experiments for U373-MG human glioma cell migration (Fig. 3A, B) [13]. The influence of confinement on migration speed is particularly significant on very stiff substrates (*K*_*e*_*/K*_0_ ≫ 1, Fig.3B). This is because confinement has a more pronounced influence on cell areal spreading and the net traction forces of polarized cells on very stiff substrates (Fig. 2B, D). On very stiff substrates, narrower channel widths lead to highly suppressed cell spreading and a very small cell aspect ratio *δ*, defined as the ratio of the shortest axis to the longest axis in the cell (Fig. 3C). The dependence of *δ* on the unitless width *W/L* is approximately linear (Fig. S7). Suppressed spreading significantly reduces the viscous drag on the nucleus as well (full line-circle in Fig. 3D). The net traction force increases linearly with decreasing aspect ratio because of the reduced radial dispersion of traction forces (Fig. 3E). In contrast, due to the limited effect of confinement on cell spreading and polarization on less stiff substrates, the reduction in the viscous drag coefficient is less pronounced (dashed diamond line, Fig. 3D), and the net traction force of polarized cells increases only marginally (Fig. 3F). This explains the varying impact of channel width on cells migrating on substrates with different stiffness. Although the above results are built upon the assumption of a well-defined cell leading edge with polarized chemical signaling, the enhancing effect of confinement on cell migration speed persists even when directional migration with persistent polarity is absent, as computed for randomly distributed initial chemical signaling (Fig. S8). Compared to persistent migration with robust polarity, cell migration under random signaling exhibits lower speeds. Notably, the increase in the mean migration speed with confinement is accompanied by an increase in the variance (Fig. S8B). This is because the net traction forces in confined channels experience larger fluctuations due to the randomness of the polarization direction (Fig. S8C).

**Figure 3:**
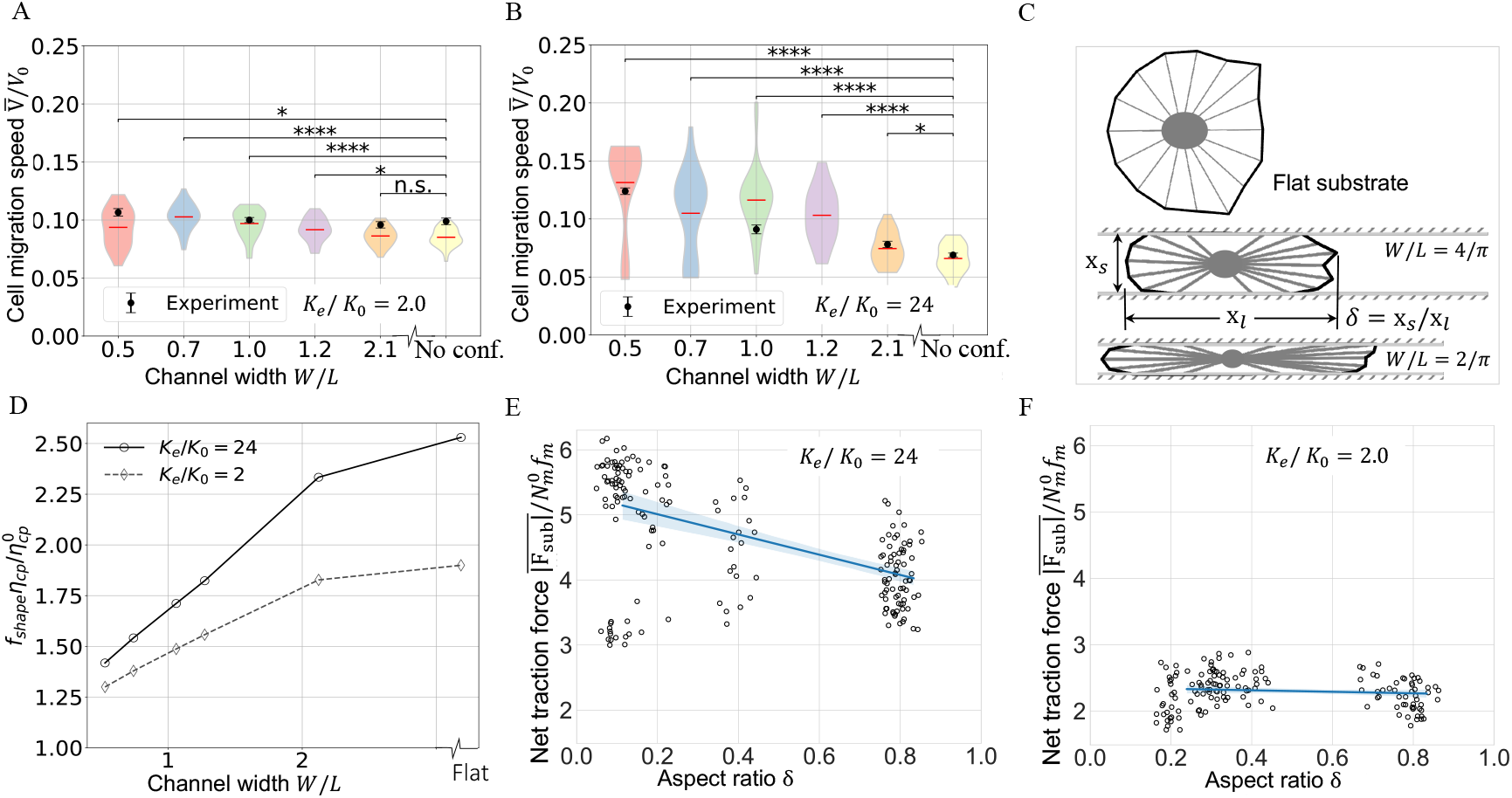
Effect of channel width on migration speed. Violin plots of 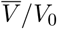 in straight channels as a function of the unitless channel width *W/L* for the substrate stiffness values (A) *K*_*e*_*/K*_0_ = 2, (B) *K*_*e*_*/K*_0_ = 24. Dots with error bars indicate the experimental data for the U373-MG human glioma cells [13]. The bar in the middle of each violin represents its arithmetic mean over more than *n >* 20 simulations (Fig. S6). Data are compared to the unitless migration speed without confinement. The p-values for statistical significance are represented by asterisks: ∗∗∗ : *p <* 0.001, ∗∗: *p <* 0.01, ∗: *p <* 0.05, and n.s.: *p >* 0.05 (Sec. S6). (C) Influence of confinement on cell polarization and morphology on very stiff substrates with *K*_*e*_*/K*_0_ = 24. The aspect ratio *δ* is defined as the ratio of the cell short axis *X*_*s*_ to its long axis *X*_*l*_. (D) Unitless viscous drag coefficient of the nucleus 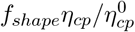 versus *W/L* for *K*_*e*_*/K*_0_ = 2 and 24. (E), (F) Unitless mean net traction force 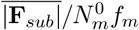 as a function of *δ* for (E) *K*_*e*_*/K*_0_ = 24 (F) *K*_*e*_*/K*_0_ = 2. In (E), (F), the lines are linear fits with a translucent 95% confidence interval band. Each dot labels single simulation result.

### Effect of channel curvature on cell spreading and migration

We next focus on the simulations of cells in curved channels to investigate the influence of curvature on cell spreading and migration speed. Our model predicts that cells exhibit reduced spreading and slower migration in channels with smaller radii or, equivalently, higher curvatures (Fig.4A, B). This can be attributed to the increase in the friction forces between the cell membrane and the wall due to the stronger contact reaction forces when a cell bends more in higher curvature channels (Movie 2). Consequently, the membrane encounters greater resistance to deformation, and the mobility of the vertices in contact with the channel walls becomes progressively hindered. The influence of curvature becomes more prominent on substrates with higher stiffness. This is because the contact length between the membrane and the wall increases with larger spreading areas on stiffer substrates, leading to higher friction forces (Movie 3). With a smaller membrane-wall dissipation coefficient *µ* (Eq. 3), we observed the reduced influence of channel curvature on cell spreading and migration speed (see Fig. S9). This verifies the critical role of the friction forces in the regulation of cell migration in curved channels.

To validate our predictions, we compared our simulations to the experiments for MDA-MB-231 cells migrating in curved channels [15]. To match the experimental setup, the simulated channels have a constant width of *W* = 8 *µm*, and two different channel radii with *R* = 50 *µm* and *R* = 100 *µm*. The substrate stiffness is taken to be *K*_*e*_ = 8 *pN/nm*, corresponding to a Young’s modulus of *E*_*sub*_ ≈ 8 *kPa* used in the experiments [15] [42]. It is evident that both the cell length and migration speed in channels with higher curvatures decrease significantly, in agreement with experiments (Fig. 4C, D).

**Figure 4:**
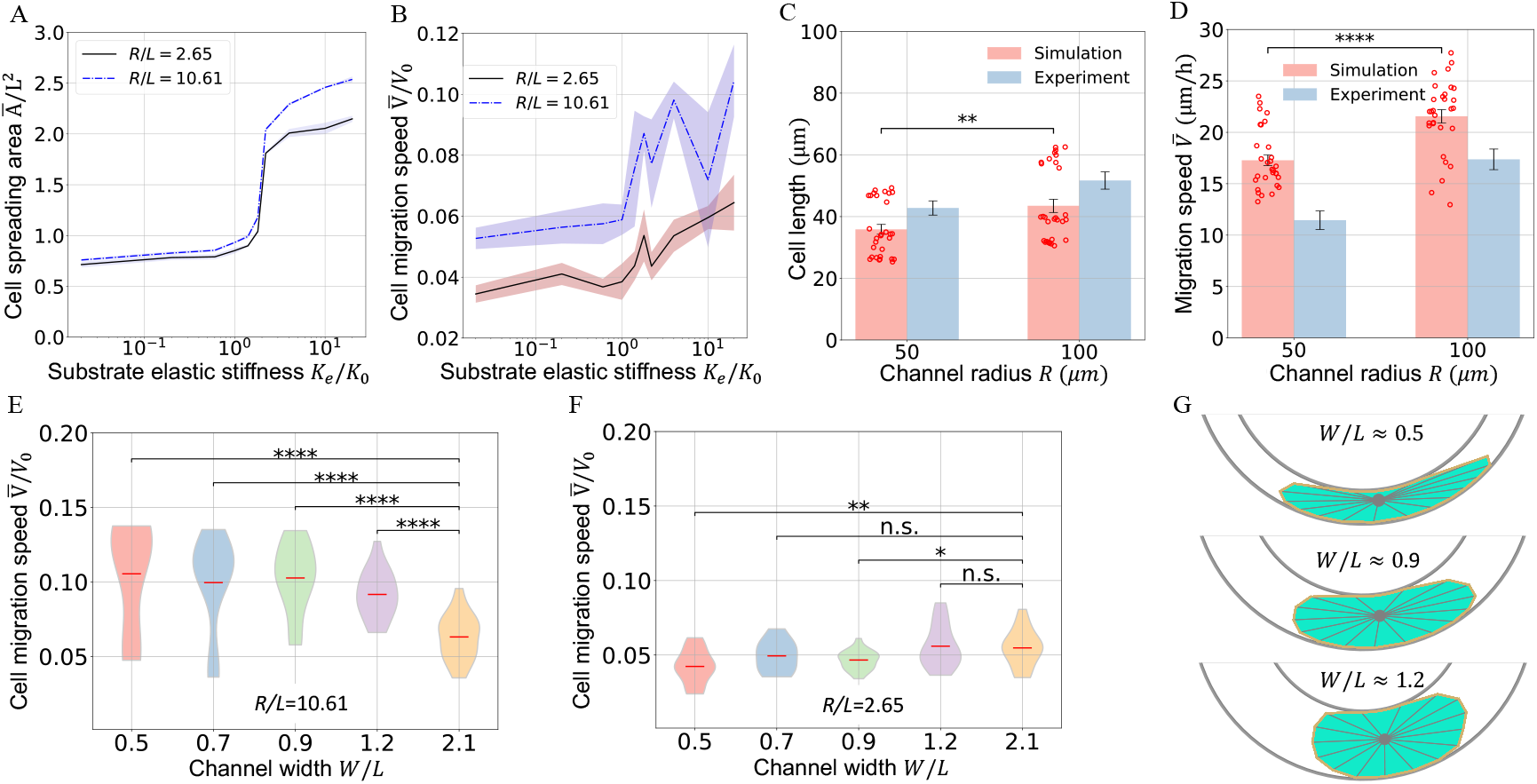
Channel curvature suppresses both spreading and migration. (A) Unitless mean spreading area *Ā/L*^2^ versus unitless substrate elastic stiffness *K*_*e*_*/K*_0_. (B) Unitless mean migration speed 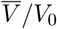 and its standard deviation as a function of *K*_*e*_*/K*_0_. The mean and standard deviation are calculated over *n* = 5 simulations at each data point. (C), (D) Comparison of our simulations with the experiments in Ref. [15] for (C) cell length and (D) migration speed in curved channels with different radii of curvature (*R* = 50 *µm* and 100 *µm*). The channels have a fixed width *W* = 8 *µm* and a fixed substrate stiffness *K*_*e*_ = 8 *pN/nm*, which correspond to the experimental parameters. Data in different channel radii are compared using t-test (*n* ≈ 20), where ∗∗ ∗∗ : *p <* 0.001, ∗ ∗: *p <* 0.01 (Sec. S6). The scatter plots display all results from simulations, with the standard deviations depicted as error bars. (E), (F) Violin plots of *V /V*_0_ versus channel width *w/L* for the channel radii (E) *R/L* = 10.61 (*R* = 200*µm*) and (F) *R/L* = 2.65 (*R* = 50*µm*), each for constant substrate stiffness *K*_*e*_*/K*_0_ = 24. The bar in each violin represents its arithmetic mean. To compare the data with wide channels (*W/L* = 2.1), a t-test was conducted with significance levels denoted by: ∗∗∗∗ : *p <* 0.001, ∗∗ : *p <* 0.01, ∗: *p <* 0.05, and n.s. (not significant) (*p >* 0.05). (G) Representative cell shapes in curved channels with different confinement widths (*R* = 50 *µm*).

To further investigate the interplay between confinement and curvature in the regulation of cell migration, we vary the channel width in curved channels. We consider a very stiff substrate (*K*_*e*_*/K*_0_ = 24) and two different unitless channel curvatures *R/L* = 10.61 and *R/L* = 2.65. For low curvature, narrower channel widths result in faster cell migration (Fig.4E), as with our prediction for straight channels (cf. Fig.3B). However, in highly curved channels (*R/L* = 2.65), increasing confinement does not promote faster migration (Fig. 4F). The observed difference can be primarily attributed to the enhanced friction forces between the cell membrane and the channel walls in more curved channels. It is evident that increasing confinement can increase friction forces due to the contact of more vertices with the walls, leading to a longer contact length of the cell membrane (Fig. 4G). In highly curved channels, which are characterized by a large friction coefficient (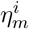 in Eq.3), cells encounter a substantial increase in the friction forces against the membrane, which largely overrides the effect of channel width and hinders migration as shown in (Fig. 4FG). In contrast, in channels with small curvature, the friction coefficient between the membrane and the channel walls remains low, similar to that in straight channels. As a result, migration speed continues to increase due to the concurrent reduction in the viscous drag force on the nucleus and the increase in the net traction force as the channel width decreases (Fig. S10).

### Combined effect of confinement geometry and matrix viscoelasticity on migration

We further sought to understand the effect of substrate viscoelasticity on cell migration in the presence of different geometric constraints, a critical factor of *in vivo* processes such as cancer invasion. For cells in straight, tightly confined channels, both cell spreading and migration dynamics depend nonlinearly on substrate viscoelasticity. In this setting, viscosity influences migration by modulating the short-term mechanical response through the instantaneous stiffness *K*_*t*→0_ = *K*_*e*_ + *K*_*a*_. Here we set the long-term substrate stiffness *K*_*e*_*/K*_0_ = 0.02 and instead varied *K*_*a*_ to modulate *K*_*t*→0_ (Fig. 5A). This approach is justified because viscoelastic effects are most pronounced when *K*_*e*_ is small; for large *K*_*e*_, *K*_*t*→0_ remains nearly constant, reducing the impact of the viscosity *γ*. On soft substrates with a low instantaneous stiffness (*K*_*t*→0_ *< K*_0_ = 5*pN/nm*), the cell spreading area has a biphasic response to the viscosity. It reaches its maximum at an optimal viscosity (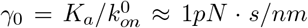 Fig. S11), where the stress relaxation timescale of the substrate *τ*_*r*_ matches the molecular clutch binding time 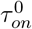 (Sec. S1). In contrast, on stiff substrates with high instantaneous stiffness (*K*_*t*→0_ *> K*_0_), the activation of adhesion reinforcement leads to enhanced cell spreading. These findings are consistent with previous studies on cell spreading dynamics on viscoelastic substrates without spatial confinement [22, 36]. Additionally, in a straight channel, a cell tends to reach its maximum migration speed on viscous substrates with a high instantaneous stiffness, where cells have large spreading areas (Fig.5A, B). Compared with cell migration on unconfined substrates, the results further demonstrate that substrate viscoelasticity has a marginal influence on cell migration in straight channels for compliant substrates but a very significant effect on substrates with high instantaneous stiffness (Fig.5C, D). The underlying mechanism can again be attributed to the close correlation between cell spreading and migration in straight channels. Confinement effectively increases the cell anisotropy when cells spread to large areas, leading to decreasing aspect ratios with increased net traction forces (Fig. S12A). An in-depth computational analysis further confirms the linear negative correlation between the cell migration speed and the aspect ratio (Fig. S12B).

**Figure 5:**
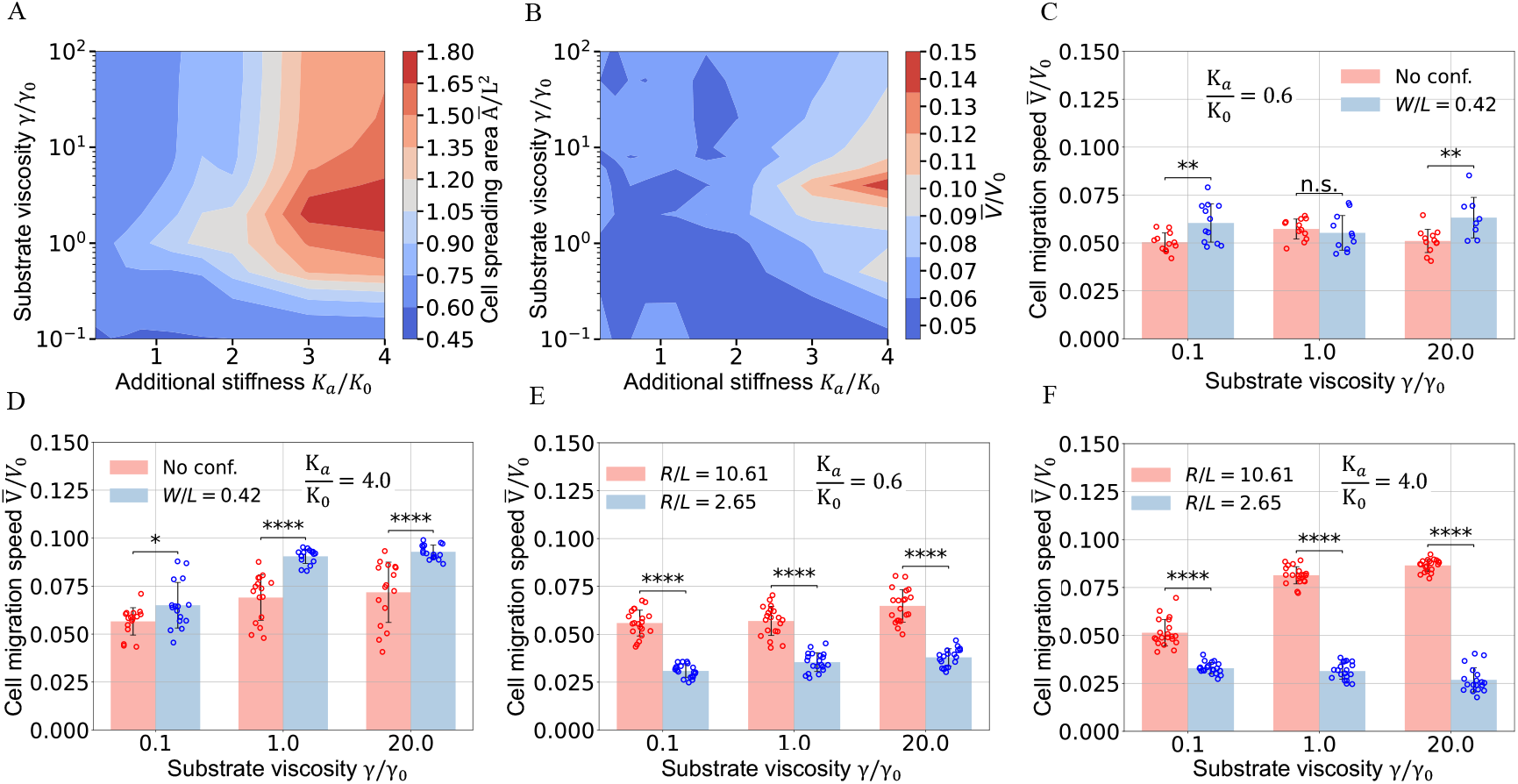
Effect of confinement and its curvature on cell spreading and migration on viscoelastic substrates. Contour plots of (A) the unitless mean cell area *Ā/L*^2^ and (B) the unitless mean migration speed 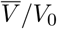 as a function of the unitless additional stiffness *K*_*a*_*/K*_0_ and unitless substrate viscosity *γ/γ*_0_ in a straight channel for a width *W/L* = 0.42 and long-term stiffness 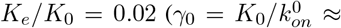 1 *pN*·*s/nm*). The contour plot contains 100 data points with *n* ≥ 5 simulations at each point. For unitless additional stiffness values (C) *K*_*a*_*/K*_0_ = 0.6, (D) *K*_*a*_*/K*_0_ = 4.0, comparison between *V /V*_0_ on unconfined substrates and in channels (*W/L* = 0.42) over different *γ/γ*_0_. For (E) *K*_*a*_*/K*_0_ = 0.6 and (F) *K*_*a*_*/K*_0_ = 4.0, comparison of 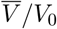 in channels with different curvatures as a function of *γ/γ*_0_. A paired t-test was performed in (C)-(F) to determine the statistical significance (*n* ≈ 20), where ∗ ∗ ∗: *p <* 0.001, ∗∗: *p <* 0.01, ∗: *p <* 0.05, and n.s.: *p >* 0.05 (Sec. S6). In (C)-(F), each dot represents one simulation result (error bars: standard deviation).

Last, we investigate the impact of channel curvature on cell migration on substrates with varying viscoelastic properties. An increase in curvature leads to a notable decrease in the cell migration speed (Fig. 5E, F), which is particularly pronounced for cells on substrates with high instantaneous stiffness and viscosity (Fig. 5F). This behavior arises from the larger friction coefficient associated with higher curvature (Eq. 3). On viscous substrates with high instantaneous stiffness, the larger spreading cells exhibit increased contact areas with the walls, resulting in amplified frictional forces (Movie 4). Moreover, in curved channels (*R/L* = 2.65), the influence of additional stiffness *K*_*a*_ and viscosity *γ* on migration is less pronounced compared to straight or low-curvature channels. For constant viscosity, increasing *K*_*a*_ leads to only a marginal decrease in migration speed, while for substrates with the same high instantaneous stiffness (*K*_*a*_*/K*_0_ = 4), increasing *γ* also leads to a marginal speed decrease. This contrasts with straight-channel results, where increasing stiffness and viscosity generally enhance migration speed. The discrepancy stems from different mechanical interactions: In straight channels, larger cell spreading boosts migration speed by increasing net traction forces while the increase in wall friction remains marginal. In highly curved channels, however, large cell spreading exacerbates wall friction due to cell bending, overriding the gain due to increased net traction. These results demonstrate that channel curvature has a more substantial impact on cell migration than the mechanical properties of the substrates alone.

## Discussion

Mesenchymal migration dynamics strongly depends on the interactions between the cell and the mechanics and interstitial geometry of the ECM. However, how these factors regulate cell migration remains unclear due to the absence of a theoretical framework that considers the contact mechanics of a migrating cell in compliance with such a complex microenvironment. To bridge this theory gap, we have developed a multiscale theory that integrates cell-matrix interactions, cell-wall contact mechanics, and intracellular chemomechanical dynamics. Our model illuminates the mechanisms governing cell migration on viscoelastic substrates in straight or curved confinement.

Our model replicates the experiments for the cell spreading area and migration speed in straight channels in Ref. [13] as a function of substrate stiffness and channel width. Our results present two key findings for constrained migration on very stiff substrates: First, strong confinement (low channel width) reduces spreading, resulting in a lower viscous drag on the nucleus. Second, anisotropic spreading enforced by the channel walls aligns the traction forces across neighboring FA sites, increasing the net traction force along the migration direction. These two factors transform the typical biphasic relationship between the cell speed and stiffness of unconfined substrates into a monotonic relationship in straight channels.

Our simulations also uncover the impact of channel curvature on cell migration. We find that cells migrate less efficiently and spread over a smaller area in channels with higher curvatures, in agreement with experiments [15]. This phenomenon arises from the augmented friction forces between the cell membrane and channel walls. As the cells migrate in curved channels, the reaction forces from the channel walls grow nonlinearly to counteract cell bending and thereby induce a higher friction at the cell-wall interface (Eq. 3). Highly curved and narrow channels substantially impede cell migration speed by merging the effects of an elevated friction coefficient with an increasing contact length. Consequently, the impact of increased confinement on migration speed is preempted by the channel curvature and even becomes detrimental. As a result, channel curvature shifts the positive correlation between the migration speed and confinement into a negative one where migration speed decreases with lower channel width.

We further explored the combined effect of confinement geometry and substrate viscoelasticity on cell spreading and migration dynamics. We find that confinement in straight channels does not alter the relationship between the cell spreading area and substrate viscoelasticity, in alignment with our previous results for migration on unconfined substrates [36]. Specifically, cells in a straight confinement attain their maximum spreading area at the same optimal viscosity with that on flat, unconfined substrates. On the other hand, a decrease in straight channel width generally enhances the migration speed, particularly on viscous substrates with high instantaneous stiffness. This effect is again attributed to the enhanced net traction force experienced by the highly polarized cell.

As opposed to straight channels, curved channels impose geometric penalties that dominate over substrate mechanics. Highly curved channels (*R/L* = 2.65) suppress migration speed by up to 50% on viscous, stiff substrates, even though such substrates enhance speed in straight channels. This is mainly because curvature increases membrane-wall friction, which outweighs the traction gain through viscous stress relaxation. These findings highlight that curvature overrides the typical speed-enhancing effects of substrate stiffness and viscosity observed in straight channels. The interplay between confinement geometry and viscoelasticity thus reveals a broader mechanical control of cell motility, where channel curvature emerges as a critical regulator with direct relevance to *in vivo* contexts. Such contexts include cancer cell metastasis through narrow, irregular tissues, where geometric cues may dictate motility more than bulk substrate mechanics alone.

While our model simplifies confinement as a 2D channels with rigid walls and uniform curvature, *in vivo* ECM microenvironments exhibit far greater geometric and mechanical complexity. First, biological channels often have locally varying curvatures, which can generate asymmetric friction forces and spatially heterogeneous resistance. Our current framework, which assumes constant curvature, can be readily extended to handle smoothly varying geometries, such as sinusoidal, parabolic, or arc-shaped channels, by defining position-dependent curvature. For highly irregular shapes where curvature is not representable by a function, further generalization using discretized geometries or image-based inputs would be required. Second, *in vivo* channel walls can deform under cell-generated forces. Softer walls may reduce the normal contact forces and tangential friction, but local deformations, such as convex 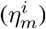 bulges formed by cellular pushing, can generate reaction forces that oppose migration. This drag can counteract reduced friction, potentially slowing the cell in softer confinement. Therefore, in straight channels, very soft walls might transform the monotonic speed-ECM stiffness relationship observed in Fig. 2B into a biphasic curve. In curved channels, such drag amplification could further reinforce the curvature-induced slowing effect. To fully capture these dynamics, incorporating viscoelastic wall mechanics would enable the model to reflect how cells respond to and actively reshape the physical constraints through bidirectional feedback between cellular forces and wall deformation.

Whereas our model captures lateral confinement on a flat substrate, real tissues impose fully 3D constraints where curvature may arise both in the wall geometry and the underlying substrate. In such microenvironments, additional curvature-dependent effects, such as traction enhancement on convex substrates or vertical reorientation along curvature ridges [43, 44], may emerge. Thus, while our core findings are likely applicable to simple 3D confinements like cylindrical tunnels, more complex 3D structures involving curved substrates may alter migration behavior through mechanisms not captured in our current 2D framework. Future extensions incorporating full 3D curvature and viscoelastic embedding could help resolve these effects. More importantly, ECM microstructure exhibits important structural features, such as matrix porosity and fiber alignment. Although channel studies offer valuable insights into cell migration within confined spaces, they cannot fully capture the complexity of the ECM fibrillar structure or long-range cell-cell force transmission mediated by ECM fiber networks (e.g., collagen or fibronectin). In such systems, forces generated by one cell propagate through aligned ECM fibers to distant cells, coupling mechanical activity across the tissue. A holistic understanding of how heterogeneous ECM structure guides cell movement will require coupling our vertex-based mechanics with finite element or network-based ECM models, as well as complementary experimental methodologies [27, 45–47].

Our multiscale model also simplifies several complex intrinsic mechanisms in cell migration. Our framework is tailored to adherent cells, where substrate traction forces and actin-based protrusion are essential drivers of movement. However, in extreme confinement (*W/L <* 0.3), mesenchymal cells can switch to an adhesion-free amoeboid phenotype where cell migration is predominantly controlled by the actin retrograde flow, and FA formation is altogether missing [12, 14]. Notably, existing models have effectively elucidated adhesion-independent migration dynamics [48–51], yet adherent cell migration under confinement remains relatively understudied. In this context, our model provides a valuable framework to further investigate how geometric and mechanical cues influence mesenchymal-to-amoeboid transition in confined environments.

Our goal here is to establish a simplified framework isolating mechanical effects, such as membrane-wall friction, curvature-induced stiffness, and nuclear drag, without the added complexity of biochemical signaling. As such, curvature-dependent behaviors in our model (e.g., cell membrane-wall friction and membrane stiffness) are treated as purely mechanical. Nevertheless, we recognize that curvature-sensitive biochemical signaling, such as cytoskeletal remodeling and localized Rho GTPase activation, could synergize with mechanical feedbacks to modulate migration. For instance, membrane curvature sensors or ion channels might alter clutch kinetics or actin polymerization rates in the motor-clutch model. Future extensions of our work could integrate mechanochemical coupling rules to explore how biochemical and mechanical signals jointly regulate migration, thereby bridging the gap between our current mechanical framework and the complexity of *in vivo* microenvironments.

## Conclusion

In conclusion, we have presented a multiscale theory for the migration of mesenchymal cells in complex microenvironments, including factors such as channel confinement, curvature, and ECM mechanical properties. Our multiscale model explains experimental spreading and migration data in straight and curved channels [13, 15]. Furthermore, we offer experimentally testable predictions for the combined influence of ECM viscoelasticity and confinement geometry on cell spreading and migration. Overall, our computational framework delivers valuable physical insights into how the ECM mechanics and geometric constraints jointly regulate mesenchymal migration.

## Supporting Information

Supplementary text, 12 figures, and a table are available in the appended PDF document.

**Movie 1**. Cell migration on elastic substrates with varying stiffness under confinement (*W/L* = 0.53 channel width).

**Movie 2**. Cell migration in curved channels with *R/L* = 2.65 and *R/L* = 10.61 on an elastic substrate with stiffness *K*_*e*_*/K*_0_ = 4.

**Movie 3**. Cell migration in highly curved channels with different substrate stiffnesses *K*_*e*_*/K*_0_ = 2 and *K*_*e*_*/K*_0_ = 20.

**Movie 4**. Cell migration in highly curved channels with different substrate viscoelasticities *γ/γ*_0_ = 0.1 and *γ/γ*_0_ = 10.

## Author Contributions

W.S.: conceptualization, formal analysis, investigation, software, validation, visualization, writing – original draft, review and editing; C.N.K.: conceptualization, formal analysis, project administration, resources, validation, writing – original draft, review and editing.

## Acknowledgments

We thank the College of Science and the Center for Mathematics of Biosystems at Virginia Tech for financial support.

## Data availability

The source codes of the simulations can be downloaded at https://github.com/nadirkaplan/confined_migration.

