## Supplemental materials for "A multiscale theory for mesenchymal cell migration in straight or curved channel confinement"

### S1 Modeling cell migration on flat substrates

Our simulations of cell migration on flat substrates without confinement are based on the whole-cell model we developed in [1]. The model effectively reproduces the spreading and migration dynamics of mesenchymal cells with adhesion reinforcement regime. Under confinement and curvature, Eqs. S8a-S8b must be modified by Eqs. 1-4 in the main text to account for the spreading dynamics of vertex in contact with the walls. In highly curved channels, Eq. S13 must be modified by Eq. 5 to accurately calculate the nucleus position. For scenarios not explicitly detailed in the main text, such as vertex spreading in microchannels without wall contact or nucleus movement in straight channels, calculations can still be carried out using the model equations originally proposed for cells on flat substrates.

#### S1.1 Equations of motion at subcellular scale

The active radial spreading of a vertex with a speed $V_{s,n}^i$ is fueled mainly by F-actin formation with a polymerization speed $V_p^i$ and counteracted by the retrograde G-actin flow with a speed $V_r^i$ , which yields

$$V_{s,n}^i = V_p^i - V_r^i \,. \tag{S1}$$

The polymerization rate at the vertex $i$ is given by the ratio of the active Rac1 concentration $R_a^i$ to its mean $\langle R_a^i \rangle$ averaged over all vertices as $V_p^i \equiv \frac{R_a^i}{\langle R_a^i \rangle} V_p^0$, where $V_p^0$ is the reference polymerization speed. The retrograde actin flow speed $V_r^i$ is promoted by the net resistance force against protrusion $F_p^i$ due to the cell membrane elasticity and myosin contractions (Fig. 1B) and impeded by an elastic restoring force $F_c^i$ due to the formation of molecular bonds by proteins such as integrins, talin, and vinculin between F-actin and ECM. We thus propose a phenomenological relation

$$V_r^i = V_0 \left( 1 - \frac{F_{am}^i}{N_m^i f_m} \right) \,, \quad F_{am}^i \equiv F_c^i - F_p^i \,, \tag{S2}$$

where $V_0$ is the unloaded myosin motor speed, $N_m^i$ is the number of active myosin motors, and $f_m$ is the force that stalls the activity of one myosin motor. Due to the increased iteration requirements for calculating $V_r^i$ when reaching its minimum during loading-unloading focal adhesion cycles, we enforce the condition $V_r = \max(V_r, 0)$ to ensure a non-negative retrograde velocity when convergence is not attained within a limited number of iterations. RhoA is known to induce myosin motor activation, leading to stress fiber formation and contractility [2, 3]. Therefore, we assume that the myosin motor number is controlled by the active RhoA concentration $\rho_a^i$ and its mean $\langle \rho_a^i \rangle$ averaged over all vertices as $N_m^i(t) \equiv \frac{\rho_a^i}{\langle \rho_a^i \rangle} N_m^0$ , where $N_m^0$ is the reference myosin motor number. In Eq. S2, the elastic restoring force $F_c^i$ is determined from the FA dynamics with adhesion reinforcement, which is triggered by the stiffness of the viscoelastic substrate. On the other hand, the protrusion force $F_p^i$ is set by the local force balance at each vertex in the presence of the nonlinear strain stiffening of the cytoskeleton. To calculate these two force strengths, we detail each of those processes next.

##### S1.1.1 Focal adhesion dynamics with adhesion reinforcement

To account for the adhesion reinforcement due to talin unfolding, we extend the augmented motor-clutch model for FA dynamics introduced in Refs. [4, 5] in order to calculate the clutch force $F_c^i$ at each vertex. By denoting average displacements of all bounded clutches at the filament end by $x_r^i(t)$ and the displacement of the substrate by $x_{sub}^i$, the engaged clutch is represented by a Hookean spring with tension $f_c^i = K_c(x_r^i - x_{sub}^i)$ ($K_c$ : spring stiffness) [6, 7]. At any instant $t$, the unbounded clutches must associate with a rate $k_{on}^i$. For the talin-low cells, a constant association rate $k_{on}^0$ is typically assumed, corresponding to a constant clutch binding timescale $\tau_{on}^0 \equiv 1/k_{on}^0$ [5, 8, 9]. We introduce the adhesion reinforcement by assuming that, when the time-averaged clutch force $\left\langle f_a^i \right\rangle_{\tau_l} \equiv \int_0^{\tau_l} f_c^i dt/\tau_l$ ($\tau_l$ : variable focal adhesion lifetime) is above a threshold force $f_{cr}$, the clutch binding rate will increase per [4, 5],

$$k_{on}^i = k_{on}^0 \left(1 + e^{\zeta(\left\langle f_a^i \right\rangle_{\tau_l} - f_{cr})}\right) , \tag{S3}$$

where $\zeta$ is a characteristic inverse force scale. Note that $\left\langle f_a^i \right\rangle_{\tau_l}$ is determined by solving an isolated motor-clutch system separately (Eqs.S1-S6), considering the updated protrusion force $F_p^i$ and the updated number of clutches and motors at vertex $i$ at time $t$. Due to its time dependence, $\left\langle f_a^i \right\rangle_{\tau_l}$ at every vertex must be updated every $N = 4000$ time step.

Once formed, the molecular clutches unbind at a dissociation rate $k_{off}^i$ that depends on the clutch tension $f_c^i$. To that end, we use the functional form $k_{off}^i \equiv k_{r0} \exp\left(f_c^i/f_{r0}\right) + k_{c0} \exp\left(-f_c^i/f_{c0}\right)$. Here, $k_{r0}$ and $k_{c0}$ denote the unloaded off-rate and the unloaded catch-rate, respectively, $f_{r0}$ is the characteristic rupture force, and $f_{c0}$ is the characteristic catch force. It follows that the fraction of the engaged clutches $(0 \leq P^i(t) \leq 1)$ is governed by the mean-field rate equation [5–7],

$$\frac{dP^i}{dt} = k_{on}^i \left(1 - P^i\right) - k_{off}^i P^i . \tag{S4}$$

Denoting the number of available clutches as $N_c^i$, the total clutch force at the vertex $i$ is then given by $F_c^i = P^i N_c^i f_c^i$. Here we incorporate the critical role of the Rac1 proteins in focal complex assembly by relating $N_c^i$ to the local Rac1 concentration, i.e., $N_c^i(t) \equiv \frac{R_a^i}{\langle R_a^i \rangle} N_c^0$, where $N_c^0$ denotes the reference clutch number [10–12]. The mechanical equilibrium condition at the cell-substrate interface demands that the total force sustained by the engaged clutches must be balanced by the substrate deformation, leading to

$$F_{sub}^i = F_c^i = P^i N_c^i K_c \left(x_r^i - x_{sub}^i\right) . \tag{S5}$$

In Eq. S5, the substrate displacement $x_{sub}^i$ is an unknown to be determined from a constitutive model for the substrate viscoelasticity, which we focus on next.

##### S1.1.2 Constitutive model for substrate viscoelasticity

Our previous implementation utilized the classical Kelvin-Voigt model for the substrate relaxation dynamics [9]. However, since the Kelvin-Voigt model predicts a very rigid nonphysical behavior when $t < \tau_r$ ($\tau_r$ : substrate relaxation timescale) [13], it can result in a premature adhesion reinforcement.

Comparatively, the standard linear solid (SLS) model allows for a better physical modeling by including a Maxwell arm (Fig. 1 B) [14]. It has also been demonstrated that the SLS model can capture the prominent relaxation timescale of the viscoelastic substrates fabricated by combining covalent and supramolecular crosslinking [5]. The SLS model expresses the constitutive relationship between $x^i_{sub}$ and the substrate force $F^i_{sub}$ as

$$(K_e + K_a)\gamma \dot{x}^i_{sub} + K_a K_e x^i_{sub} = K_a F^i_{sub} + \gamma \dot{F}^i_{sub}\,, \tag{S6}$$

where $K_e$ is the elastic stiffness at $t \to \infty$, $\gamma$ is the substrate viscosity, and $K_a$ is the additional stiffness that governs the substrate relaxation with a timescale $\tau_r \equiv \gamma / K_a$ . A stress relaxation test from a constant strain yields the instantaneous and long-term stiffness of the substrate as $K_{t\to 0} = K_e + K_a$ and $K_{t\to\infty} = K_e$ , respectively. The instantaneous stiffness $K_{t\to 0}$ characterizes the initial elastic response of the substrate when the viscous deformation and stress relaxation have not yet taken place in the limit $t \to 0$. Thus, cells with a very short focal adhesion lifetime ($\tau_l < \tau_r$) can only sense the instantaneous stiffness. In contrast, the long-term stiffness $K_{t\to\infty}$ refers to the residual substrate stiffness after the viscous stresses have relaxed. For an elastic, soft substrates, the clutch force of an individual bond $f^i_c$ develops slowly with a long lifetime $\tau_l \approx \frac{N^0_m f_m}{V_0 K_e}$ [1], contributing to a low time-averaged clutch force $\left\langle f^i_a \right\rangle_{\tau_l} < f_{cr}$. Consequently, the clutches associate with a constant binding rate $k^i_{on} = k^0_{on}$, corresponding to a constant binding timescale $\tau^0_{on} = 1/k^0_{on}$. The clutch lifetime decreases as the substrate stiffness $K_e$ increases. When $\tau_l \leq \tau^0_{on}$, the limited lifetime leads to the insufficient formation of the bounded clutches and thus the force shared by an individual bond increases $\left\langle f^i_a \right\rangle_{\tau_l} \geq f_{cr}$, triggering the adhesion reinforcement regime. Therefore, The equality $\tau_l = \tau^0_{on}$ defines a threshold stiffness $K_0 = N^0_m f_m k^0_{on} / V0$, which allows us to classify a substrate as "stiff" when $K_e > K0$ and "soft" when $K_e < K_0$.

##### S1.1.3 Cytoskeletal stiffening

The passively deforming cell cytoskeleton, which consists of microtubules and intermediate filaments, is represented by multiple springs in our model (Fig. 1 A). Here we will assume that these cytoskeletal "springs" undergo strain-stiffening during large spreading events and thus exhibit nonlinear elasticity. This assumption is backed by the experiments performed, e.g., on NIH-3T3 fibroblasts that reveal a strong correlation between the cell rigidity and cell area during large spreading events [15]. Assuming this behavior is mechanically driven and thus must be generic across animal cells, we propose a phenomenological equation for the cytoskeletal stiffness $K_{cs}$ in terms of the cell area $A$: Denoting the position vector of the cell nucleus by $\mathbf{x}_{nuc}$ and defining the length of a cytoskeletal spring as $r^i \equiv |(\mathbf{x}_{nuc} - \mathbf{x}^i)|$ , the cytoskeletal restoring force in our model is governed by the differential equation

$$\frac{dF^i_{cs}}{dr^i} = K_{cs}(A) = K^b_{cs} + \Delta K_{cs} e^{\beta A}\,, \tag{S7}$$

where $K^b_{cs}$ denotes the baseline stiffness. The term $\Delta K_{cs} e^{\beta A}$ is introduced to describe the exponential hyperelastic behavior [16, 17], with the values of $\Delta K_{cs}$ and $\beta$ obtained through linear regression of the experimental data in [15].

##### S1.1.4 Local mechanical equilibrium

We enforce local force balance at each vertex to determine the radial protrusion force $F_p^i$ and the polar spreading speed $V_{s,\tau}^i$. The cell membrane possesses a notable mechanical rigidity that enables it to endure a variety of stresses, which is vital for maintaining the integrity of cell structures [18–21]. We model it as a closed loop of Hookean springs with stiffness $K_m$ between neighboring vertices. The tensile forces acting on the $i^{th}$ vertex can be expressed as $\mathbf{F}_{m,\pm}^i = K_m(\mathbf{l}_\pm^i - \mathbf{l}_0)$, where $\mathbf{l}_+^i$ and $\mathbf{l}_-^i$ are the distance vectors between vertex $i$ and its two neighbors, and $\mathbf{l}_0$ is the initial distance between two adjacent vertices. The drag force on vertex $i$, resulting from hydraulic resistance in the extracellular medium, can be expressed as $F_\eta^i = \eta_m^i l^i \mathbf{V}_s^i$, where $l^i \equiv (|\mathbf{l}_+^i| + |\mathbf{l}_-^i|)/2$ defines the average membrane length about the $i^{th}$ vertex. The drag coefficient $\eta_m^i$ is proportional to the fluid viscosity but inversely proportional to the matrix permeability [22–25]. In addition to the membrane forces, each vertex experiences a protrusion force $F_p^i$ and a cytoskeletal radial force $F_{cs}^i$. Thus, the net force balance at each vertex in the radial direction $\hat{\mathbf{n}}^\mathbf{i}$ and the polar direction $\hat{\boldsymbol{\tau}}^i$ is given by

$$F_p^i - F_{cs}^i + \left(\mathbf{F}_{m,+}^i + \mathbf{F}_{m,-}^i\right) \cdot \hat{\mathbf{n}}^\mathbf{i} - \eta_m l^i V_{s,n}^i = 0\,, \tag{S8a}$$

$$\left(\mathbf{F}_{m,+}^i + \mathbf{F}_{m,-}^i\right) \cdot \hat{\boldsymbol{\tau}}^i - \eta_m l^i V_{s,\tau}^i + \left(F_{rp,+}^i + F_{rp,-}^i\right) = 0\,. \tag{S8b}$$

where the last term $F_{rp,\pm}^i$ on the left-hand side of Eq. S8b represents repulsive forces in polar direction. These forces only become non-zero when the cytoskeletal elements of adjacent vertices almost overlap within highly curved channels (see Sec. S4). Eq. S8a and S8b yield $F_p^i$ and $V_{s,\tau}^i$, respectively. Altogether, Eq. S1–S8b fully determine the vertex spreading velocities $\mathbf{V}_s^i$ when the nucleus displacement $\mathbf{x}_{nuc}$ and the dynamical GTPase concentrations at each vertex are computed at the cellular scale.

#### S1.2 Equations of motion at cellular scale

Next, we explain the global mechanical equilibrium that governs the nucleus motion and the intracellular Rho-GTPase dynamics, which directly influence the aforementioned subcellular processes.

##### S1.2.1 Viscous drag on deformed nucleus

Our model quantifies cell translocation by the net translation of the cell nucleus, which balances the forces between the cytoskeletal microtubules and intermediate filament bundles. At the cell periphery, these cytoskeletal complexes are linked to the F-actin at the FA sites, transmitting the net traction force from the extracellular matrix ($F_{sub}^i$) and the protrusion force from the cell boundary ($F_p^i$) to the cell nucleus (Fig. 1 A). This tight linkage to the rest of the cell can deform the nucleus when the cell flattens under large spreading on stiff substrates [26]. To balance these peripheral and cytoskeletal forces, the nucleus undergoes viscous drag within the cytoplasm while it deforms under cell flattening. The viscous drag force on a particle is commonly described by the product of a drag coefficient, a characteristic particle size, and the velocity of the particle relative to the surrounding medium. Although the cytoplasm as the surrounding medium may undergo convective flows due to the intracellular biomolecular and organelle dynamics, its velocity around the nucleus can point in any direction at a certain time instant, either facilitating or opposing the movement of the nucleus at that instant. Thus, over extended periods of time, the contribution of the cytoplasmic flow to the motion of the nucleus can be neglected, and we treat the cytoplasm as a stationary viscous medium for the calculation of the nucleus velocity [1].

In the case of a deformed nucleus with irregularities, we incorporate a shape factor denoted as $f_{shape}$ into the analysis [27]. This leads to the following expression for the drag force:

$$\mathbf{F}^{\eta}_{nuc} = -f_{shape}\mathcal{L}_{nuc}\eta_{cp}\dot{\mathbf{x}}_{nuc}\,, \tag{S9}$$

where $\eta_{cp}$ represents the viscosity of the cytoplasm and $\mathcal{L}_{nuc}$ is the characteristic particle size. For simplicity, we take $\mathcal{L}_{nuc} \equiv 6\pi r^0_{nuc}$ corresponding to Stokes' flow as a first-order approximation to the cytoplasmic domain within finite cell height ($r^0_{nuc}$: the initial radius of the nucleus). The shape factor $f_{shape}$ quantifies the deviation of the nucleus from a perfect sphere with $f_{shape} = 1$ . The Corey shape function can be used to determine $f_{shape}$ based on the nucleus's three principal lengths [27, 28],

$$f_{shape} = \left(\frac{l_{max}l_{med}}{l^2_{min}}\right)^{\alpha}\,, \tag{S10}$$

where the exponent $\alpha = 0.09$ was obtained by fitting the experimental drag coefficient of non-spherical particles under Stokes flow [27]. For the deformed nucleus, the aspect ratio defined by the longest and the shortest dimensions can be related to the cell spreading area by $\delta \equiv \frac{l_{max}}{l_{min}} = \chi A + 1$ , where the exponent $\chi = 0.0024$ is obtained by fitting the experimental cell shape data in Ref. [29]. The intermediate dimension $l_{med}$ corresponds to the length of the minor axis of the nucleus on the $x - y$ plane, and it was found to have a constant ratio to the length of the major axis in experiments ($l_{med} = 0.8 l_{max}$) [26]. Consequently, the nucleus shape factor can be simply related to the cell spreading area $A$ by,

$$f_{shape} = 0.98\,(\chi A + 1)^{2\alpha}\,. \tag{S11}$$

Experiments have demonstrated that the cytoplasm viscosity, like the strain stiffening, is strongly correlated with the cell spreading area [30, 31]. These experiments also indicate that the cell viscosity and cell stiffness exhibit the same trend with increasing substrate stiffness. Therefore, we assume an area-dependent cytoplasm viscosity (similar to that in Eq. S7) as

$$\eta_{cp}(A) = \eta^0_{cp} + \Delta\eta_{cp}e^{\beta A}\,, \tag{S12}$$

where $\eta^0_{cp}$ is the cytoplasm viscosity on soft substrates [32] and $\Delta\eta_{cp}$ controls the rate of viscosity increase with the cell spreading area $A$ .

At the frame of the nucleus, the condition for the mechanical equilibrium between Eq. S9, the cytoskeletal and the subcellular forces

$$\sum_{i=1}^{N}\left(F^i_{sub} + F^i_{cs} - F^i_p\right)\hat{\mathbf{n}}^i - \mathbf{F}^{\eta}_{nuc} = 0 \tag{S13}$$

yields the nucleus migration velocity, $\dot{\mathbf{x}}_{nuc}$ , equivalent to the cell velocity in our model.

##### S1.2.2 Reaction-diffusion dynamics of GTPase concentrations

As with our previous work [9], here we adopt the biochemical reaction-diffusion equations introduced in [33, 34] to describe the dynamics of active and inactive GTPases. Since the active forms of the GTPases are predominantly associated with the cell membrane, which also serves as a major site for the conversion between active and inactive forms, we track the volume fractions of the signaling proteins in three forms [35, 36]: the active membrane-bound form $(G_a^i(t) \equiv \{R_a^i(t)\,, \rho_a^i(t)\})$, the inactive membrane-bound form $(G_{in}^i(t) \equiv \{R_{in}^i(t)\,, \rho_{in}^i(t)\})$ , and the inactive cytosolic form $(G_{cp}(t) \equiv \{R_{cp}(t)\,, \rho_{cp}(t)\})$ . Given the rapid diffusion of inactive proteins in the cytosol, we assume $G_{cp}(t)$ remains uniformly distributed in the cytosol at all times. Both active and inactive membrane-bound forms diffuse with a diffusion constant $D$ across the vertices. The corresponding diffusive fluxes are given by the Fick's law in a finite difference formulation as

$$J_y^i \equiv -D\left(\frac{G_y^{i+1}/l^{i+1} - G_y^i/l^i}{|\mathbf{l}_+^i|}\right) . \tag{S14}$$

Eq. S14 takes into account the effect of the deformed cell shape on the diffusive flux by updating the vertex coordinates at each time step. Different forms of the proteins on a vertex are interconvertible with the inactive-to-active rates $A_G^i$, active-to-inactive rates $I_G^i$, and inactive-to-cytosolic association and disassociation rates $M_G^+$ , $M_G^-$ (Fig. 1 A). Altogether, the reaction-diffusion kinetics is governed by

$$\begin{cases} \dot{G}_a^i = A_G^i G_{in}^i - I_G^i G_a^i + \left(J_a^{i-1} - J_a^i\right) \,, \\ \dot{G}_{in}^i = -A_G^i G_{in}^i + I_G^i G_a^i + \left(J_{in}^{i-1} - J_{in}^i\right) + \dfrac{M_G^+ G_{cp}}{N} - M_G^- G_{in}^i \,, \\ \dot{G}_{cp} = -M_G^+ G_{cp} + \displaystyle\sum_{i=1}^{N} M_G^- G_{in}^i \,. \end{cases} \tag{S15}$$

The active Rac1 and RhoA GTPase volume fractions $R_a^i$ , $\rho_a^i$ regulate the cell migration by controlling the actin polymerization speed $V_p^i$ (Eq. S1), the number of myosin motors $N_m^i$ (Eq. S2), and the clutch number $N_c^i$ at each FA site (Eq. S5).

Definitions of rates $A_G^i$ in Eq.S15 accommodate autoactivation and antagonistic effects as well as the feedback from the mechanical deformations of the cell membrane. The RhoA activation rate $A_\rho^i$ at the $i^{th}$ vertex is composed of three terms,

$$A_\rho^i = \frac{\alpha_\rho {\rho_a^i}^3}{\left({\rho_0}^3 + {\rho_a^i}^3\right)} + \frac{\beta_\rho {R_0}^3}{\left({R_0}^3 + {R_a^i}^3\right)} + e^{-\mathcal{H}(-\Delta\theta - c_0\theta_0)\frac{\Delta\theta}{\theta_0}}\kappa_b^+ \,, \tag{S16}$$

where $R_0$ and $\rho_0$ are reference levels of the active Rac1 and RhoA on the vertex, respectively. The first term takes into account positive feedback from the active RhoA itself. The magnitude of the autoactivation effect is represented by $\alpha_\rho$ [37, 38]. The second term describes the mutual inhibition effect between Rac1 and RhoA with a rate $\beta_\rho$ [37, 38]. The last term couples the active level of RhoA to the mechanical deformations of the cell at the $i^{th}$ vertex. $\mathcal{H}()$ denotes the Heaviside step function. The initial angle at a cell vertex is labeled by $\theta_0$ , and $\Delta\theta \equiv \theta - \theta_0$ denotes the change in angle at the vertex (Fig. 1A). Here we assume that the active rate of RhoA increases when the vertex angle satisfies $\Delta\theta < -c_0\theta_0$ due to the cell polarization.

Similarly, Rac1 activation rate $A_R^i$ at the $i^{th}$ vertex also consists of three terms,

$$A_R^i = \frac{\alpha_R {R_a^i}^3}{\left({R_0}^3 + {R_a^i}^3\right)} + \frac{\beta_R {\rho_0}^3}{\left({\rho_0}^3 + {\rho_a^i}^3\right)} + e^{\mathcal{H}(\Delta\theta - c_0\theta_0)\frac{\Delta\theta}{\theta_0}} K_b^+ . \tag{S17}$$

We assume that the active rate of Rac1 soars once the increase of the vertex angle exceeds the threshold $c_0\theta_0$ due to the cell contraction. Coupling Rac1 and RhoA to mechanical deformations enables recurrent polarization and the recovery of cell deformations. Specifically, an excessive increase in $\theta^i$ at the retracting cell rear builds up $R_a^i$ to increase actin polymerization and thus reverses the retraction at the end of each migration step. A large decrease in $\theta^i$ at the protruding cell front boosts $\rho_a$ that increases actomyosin contraction to resist further protrusion.

Since the filopodial protrusions of a mesenchymal cell grow and shrink at timescales comparable to the migration times, this chemo-mechanical feedback prevents the Rac1 and RhoA dynamics from reaching a steady bistable polarized state. Instead, when the transient chemical polarity ceases, our simulation algorithm reinstates the polarization stochastically to sustain the random migration patterns in many mesenchymal phenotypes [39, 40].

### S2 Key parameters fitting based on experimental data

The fitting parameters $\epsilon$ and $\mu$ were determined by comparing numerical predictions with experimental data. In each case, the experimental result was treated as a reference value, shown as a horizontal dashed line with a shaded region indicating variability. However, it is important to note that these experimental values are independent of the fitting parameters themselves.

For $\epsilon$, the numerical predictions of migration speed were obtained for different values of $\epsilon$ and compared against experimental measurements. The predicted migration speed from simulations varied with $\epsilon$, allowing us to assess how well each choice of $\epsilon$ aligned with the experimental result (see Fig.S1). Similarly, for $\mu$, simulations were performed to obtain the predicted cell length as a function of $\mu$. The best-fit value of $\mu$ was chosen by comparing the numerical predictions against the experimentally observed cell length (see Fig.S3).

### S3 Increase of contact forces on cells in more curved channels

The contact forces between cells and channels are directly associated with the friction forces on the cell membrane, which regulate the deformation of the membrane and the movement of adjacent vertices in contact with the channel wall. Understanding how these contact forces change in channels with varying curvatures is crucial for investigating cell migration in curved environments (Fig.S2A). However, our vertex-based model, which represents the cell as a cable-truss structure without bending stiffness, cannot accurately predict the development of contact forces due to cell bending deformations in curved channels.

To address this limitation, we introduce a 2D deep beam representation for the cell, with the length to height ratio of 7.5, to capture bending deformations in confined channels. Finite element analysis is employed to analyze the changes in channel reaction forces acting on the cell under different degrees of bending. In our model, we consider an incompressible material with Young's modulus of $E = 10\ kPa$, which is a typical cytoskeleton stiffness of spreading cells. The beam is discretized using quadratic plane stress solid elements with a mesh size of $1.5\mu m \times 1\mu m$. The upper and bottom edges of the beam are constrained to mimic the channel confinement (Fig.S2B). By applying controlled displacements, we deform the beam to simulate bending shapes similar to those experienced by cells in channels with different curvatures.

As the curvature increases, the internal stress in the beam significantly rises (Fig. S2B). We can describe the average boundary traction at the contact region of the upper boundary in relation to the Cauchy stress tensor. Denoting the boundary traction as $\mathbf{T}(\mathbf{x})$ and the unit normal vector of the boundary as $\mathbf{n}$, the average boundary traction can be expressed as follows:

$$\overline{\mathbf{T}} = \frac{\int_{S_\sigma} \sigma \cdot \hat{\mathbf{n}}\, dS}{S} \tag{S18}$$

According to Newton's third law, the boundary traction forces are equal in magnitude and opposite in direction to the reaction forces exerted by the constraints on the beam. These reaction forces play a similar role to the contact forces experienced by bending cells in curved channels. Building upon this understanding, we approximate the contact force per unit length by,

$$\overline{q}_R = \sqrt{\overline{T}_x^2 + \overline{T}_y^2} \tag{S19}$$

where $\overline{T}_x$ and $\overline{T}_y$ represent the average traction ($\overline{\mathbf{T}}$) components in the x and y directions, respectively. This approximation allows us to analyze how the contact force varies with channel curvatures. Further analyses contribute to a quadratic relation between the contact force ($\overline{q}_R$) and the curvature (Fig.S2C). These contact forces ultimately influence cell migration by altering the friction forces on the cell membrane. Thus, in our vertex-based model, we incorporate curvature-dependent friction coefficients and membrane stiffness in the contact regions (as described by Equations 6 and 7 in the main text).

### S4 Calculation of the repulsive forces

A pair of repulsive forces were introduced to maintain the mechanical integrity of the cell and model stability in computation. Due to the influence of the highly curved channel on vertex polar movements, the cytoskeletal elements of the vertices located near the channel walls may get overlap or intersect. To prevent such nonphysical penetration and avoid the convergence issue, a pair of repulsive forces $\mathbf{F}^i_{rp,\pm}$ in the polar direction are added to Eq. 4b and Eq. S8b for updating vertex polar spreading velocities in highly curved channels. $\mathbf{F}^i_{rp,\pm}$ are non-zeros only when the angle between two adjacent cytoskeletal elements falls below a threshold ($\phi_i < \phi_0$ in Fig. 1C). Based on the penalty method in contact mechanics, the repulsive forces increase with the violation of the angle threshold. Here, we assume that the magnitude of the repulsive forces is proportional to the difference between the angle of the adjacent cytoskeletal elements ($\phi_i$) and the threshold value ($\phi_0$):

$$\mathbf{F}^i_{rp,+} = -F^0_{rp}\{1 - \phi_i/\phi_0\}\boldsymbol{\tau}_i \quad \text{and} \quad \mathbf{F}^i_{rp,-} = F^0_{rp}\{1 - \phi_{i-1}/\phi_0\}\boldsymbol{\tau}_i\,, \tag{S20}$$

where $\{x\} \equiv \max(x, 0)$ , and the reference penalty force is $F^0_{rp} = 10^3\ pN$.

#### S5 Initial conditions of Rho GTPases

Since chemical signaling drives cell polarization followed by directional locomotion on uniform substrates, a nonuniform initial distribution of the active Rac1 protein volume fraction $R^a_i$ is enforced with a higher value at the cell front. Likewise, a polarized initial distribution of the active RhoA protein volume fractions $\rho^a_i$ is taken with accumulation at the cell rear (Fig. S4). In contrast, the initial conditions for $R^i_{in}$, $\rho^i_{in}$ are uniform at time $t = 0$, and the cytoplasmic form $R_{cp}$, $\rho_{cp}$ can be determined from the total volume fraction conservation. When the cytoplasmic inactive Rac1 volume fraction $R_{cp}$ reaches a steady state, we reinitialize $R^i_a$ and $\rho^i_a$ stochastically to mimic the random nature of the protrusion formations in the mesenchymal cells [9].

#### S6 Statistical analysis and visualization

We performed statistical analyses using scipy.stats package in Python. Unless otherwise stated, all statistical comparisons were based on at least $n = 5$ simulations, and p-values were calculated using t-tests for comparison of results corresponding to different biophysical properties. Statistical significance, denoted by asterisks ($*p < 0.05$, $**p < 0.01$, $***p < 0.001$, $****p < 0.0001$), is a crucial measure used to determine whether observed differences between groups or conditions are meaningful or merely due to chance. When a p-value is below a certain threshold, indicated by the corresponding number of asterisks, it suggests that the observed differences are unlikely to have occurred by chance alone. In other words, there is evidence to support the presence of a meaningful difference between the groups being compared. In contrast, we use the term "n.s." (not significant) to indicate that the observed differences are not statistically significant, suggesting that they may have arisen due to random chance or sampling variability.

Various types of visual representations, including scatters, line plots, bar graphs, violin plots, and heat maps, were employed as appropriate for the respective analyses. In a violin plot, each category or group is represented by a "violin" shape, where the width of the shape indicates the density or frequency of data points at different values (see Fig. S7). The thicker sections of the violin represent regions with higher data point concentration, while the thinner sections indicate lower density.

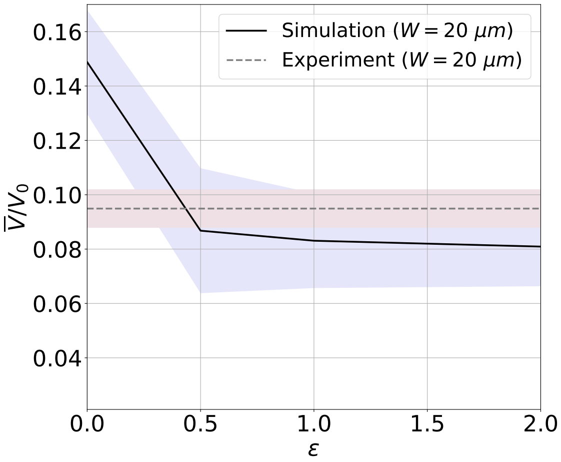


Figure S1: **The process of determining the fitting parameter $\epsilon$ by comparing numerical predictions with experimental data.** The experimental data from [41], shown as a horizontal dashed line with a shaded region, represents the target migration speed. The elastic substrate has a constant stiffness $K_e = 10\, pN/nm$ . The solid line represents the predicted migration speed from simulations as a function of $\epsilon$ , with its shaded area indicating the mean and standard deviations of numerical predictions. Each data point is based on $n \geq 5$ sets of simulations. Note that the experimental result is plotted solely for comparison and is independent of $\epsilon$.

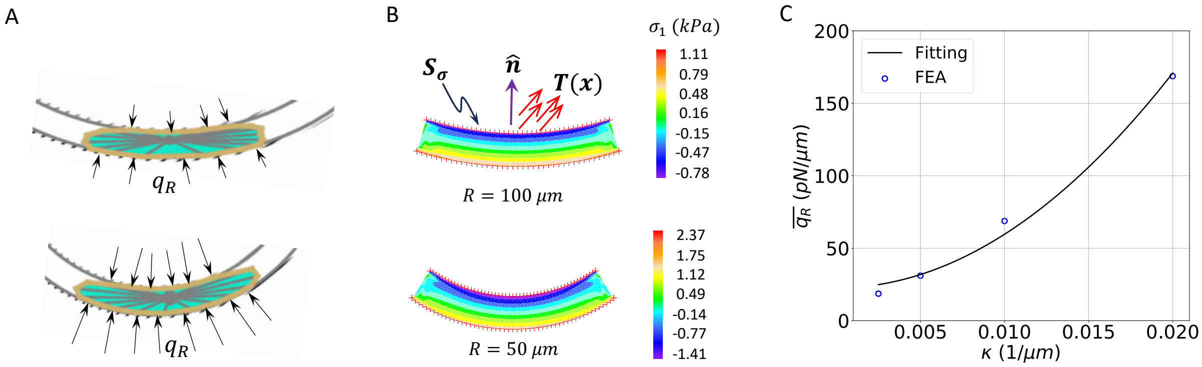


Figure S2: **Channel reaction force on a cell changes when the channel curvature varies.** (A) Depiction of elongated and bending cells within channels of varying curvatures. Arrows indicate reaction forces exerted by the channel on the cell, resisting bending deformation. (B) Stress contours of bending beams representing cells in channels with different curvatures. The symbol $S_\sigma$ represents the contact boundary under consideration. The boundary traction force at a specific point $\mathbf{x}$ is denoted by $\mathbf{T}(\mathbf{x})$. The unit normal vector of the boundary is defined as $\mathbf{n}$. (C) Relationship between channel curvatures and the corresponding reaction force per unit length at the boundaries. The data was fitted using a quadratic function given by $\overline{q}_R = a\kappa^2 + b$, where $a$ and $b$ are the fitting parameters.

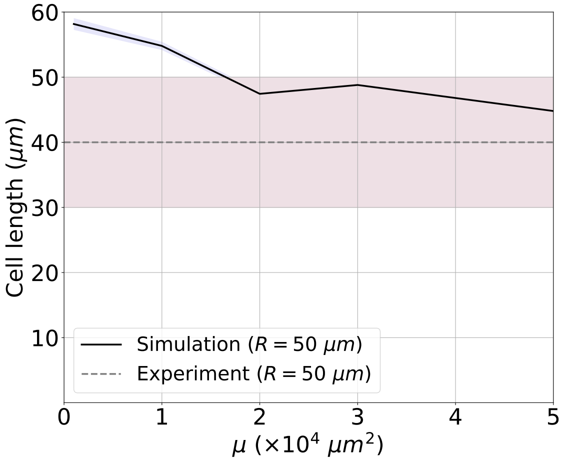


Figure S3: **The process of determining the fitting parameter $\mu$ by comparing numerical predictions with experimental data.** The experimental data of MDA-MB-231 cells in the curved and confined channel ($R = 50\,\mu m$ and $W = 8\,\mu m$), shown as a horizontal dashed line with a shaded region, represents the target cell length [42]. The solid line represents the predicted cell length from simulations as a function of $\mu$, with its shaded area indicating the mean and standard deviations of numerical predictions. Each data point is based on $n \geq 5$ sets of simulations with constant substrate stiffness, $K_e = 10\,pN/nm$. Note that the experimental result is plotted solely for comparison and is independent of $\mu$.

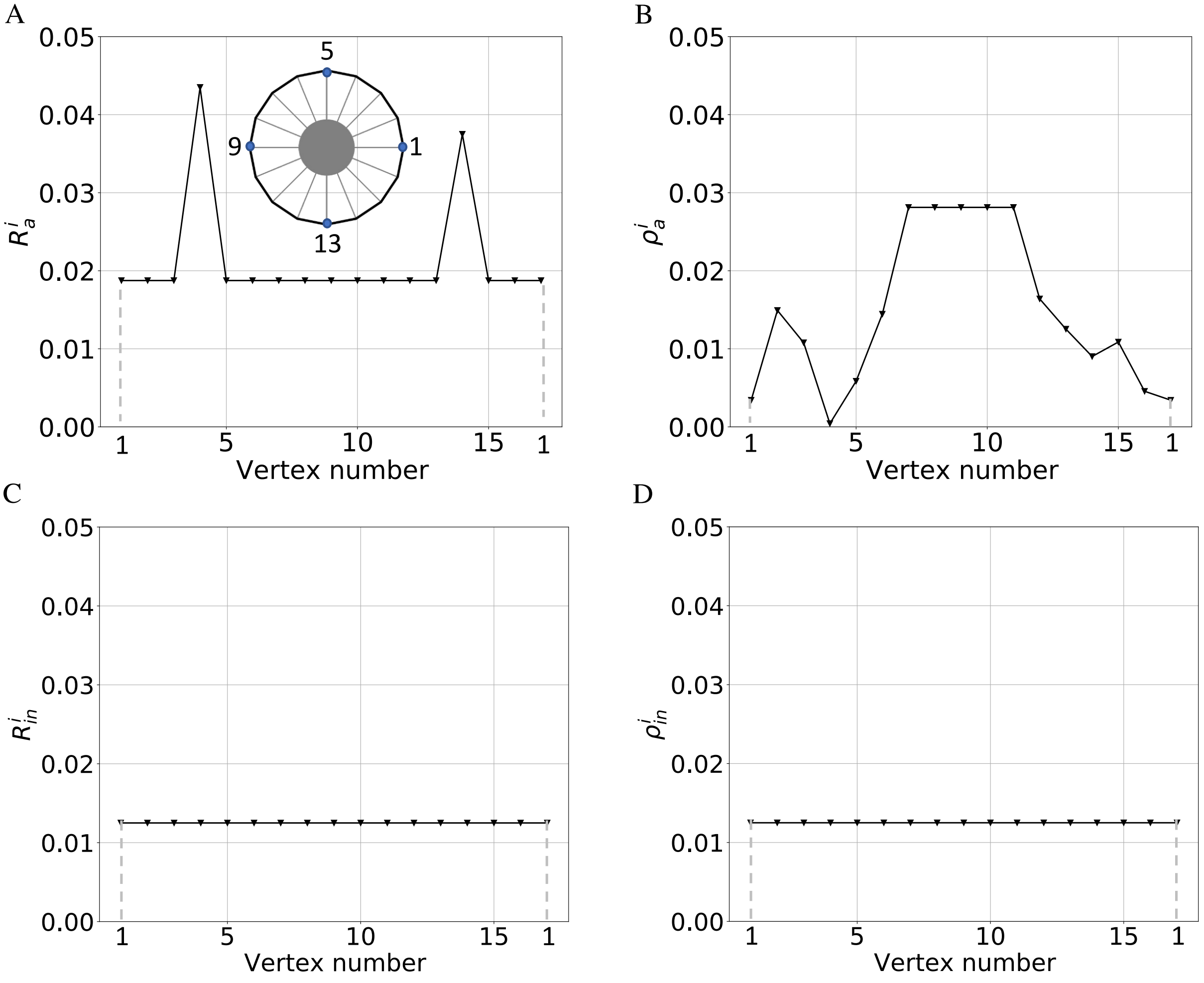


Figure S4: Initial conditions of the membrane-bound Rac1-RhoA signals for the directional migration. (A) The representative initial condition of the active Rac1 signal $R_a^i$ across all vertices ($N = 16$). Cell shape at $t = 0\ s$ illustrates the numbering of vertices. (B) Polarized distribution of the active RhoA signaling $\rho_a^i$ at $t = 0\ s$ with the uniform high concentration at the leftmost five vertices. (C-D) All other types of membrane-bound proteins, i.e., $\rho_a^i$, $R_{in}^i$, $\rho_{in}^i$, are uniformly distributed on vertices. Since the total concentration of Rac1 and RhoA are each conserved, the initial cytoplasmic concentration $G_{cp}(t = 0)$ , $G \equiv R$ , $\rho$ can be determined by $G_{cp} = 1 - \sum_{i=1}^{N} G_a^i - \sum_{i=1}^{N} G_{in}^i$ .

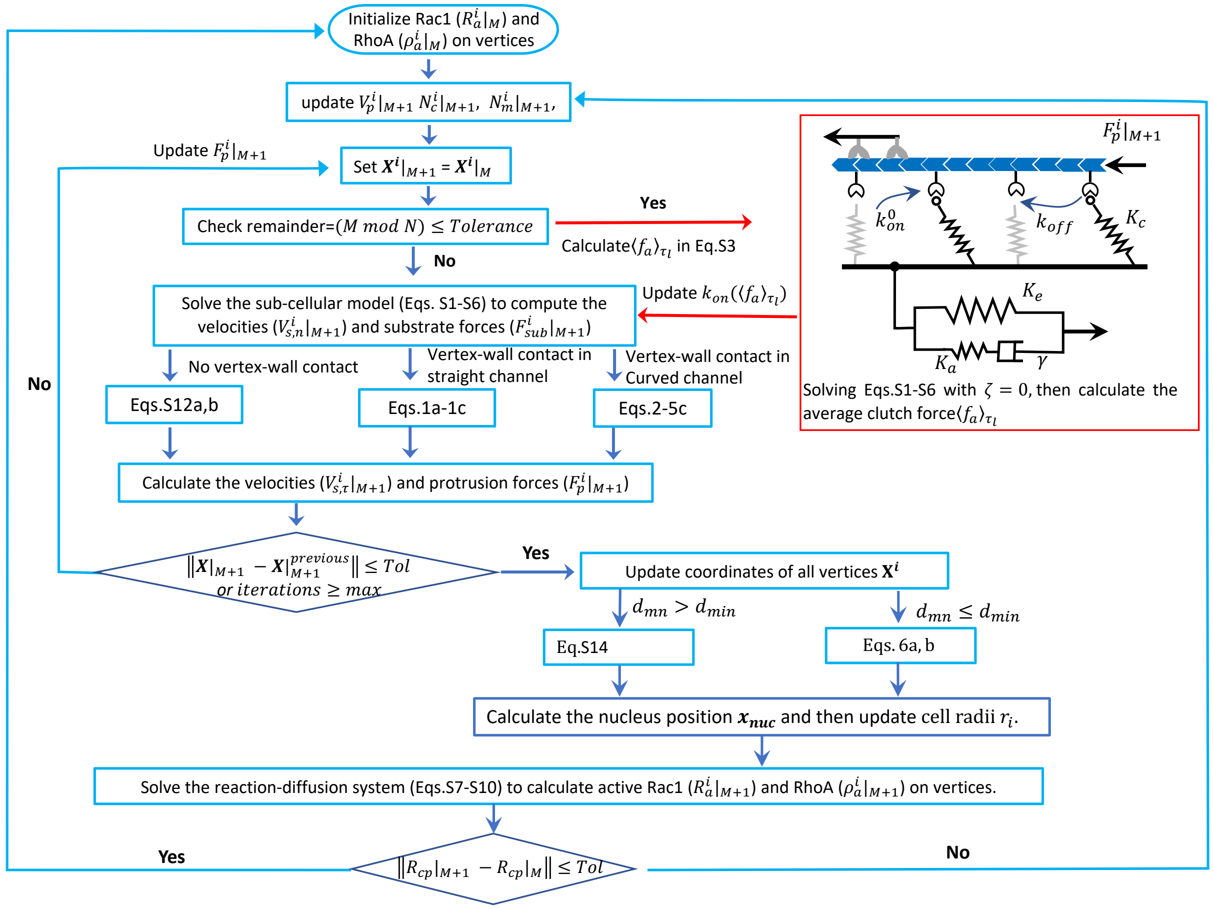


Figure S5: Flowchart of the proposed model implementation. The adhesion reinforcement condition ($\langle f_a^i\rangle_{\tau_l} > f_{cr}$) is checked every $N$ steps. At each time step $(M+1)$, cell radial spreading speed ($V_p^i|_{M+1}$), the number of clutches ($N_c^i|_{M+1}$), and the number of myosin motors ($N_m^i|_{M+1}$) at vertex $i$ are recorded. The position of vertex $i$ at time step $M$ is denoted as $\mathbf{X}^i|_M$. At time step $(M+1)$, radial ($V_{s,n}^i|_{M+1}$) and polar ($V_{s,\tau}^i|_{M+1}$) velocities of vertex $i$ are calculated, as well as the protrusion force ($F_p^i|_{M+1}$) and substrate traction force ($F_{sub}^i|_{M+1}$) of the vertex. $d_{mn}$ denotes the distance between nucleus and cell membrane in Fig.1E. A numerical tolerance of $Tol = 10^{-6}$ is set for steady state check.

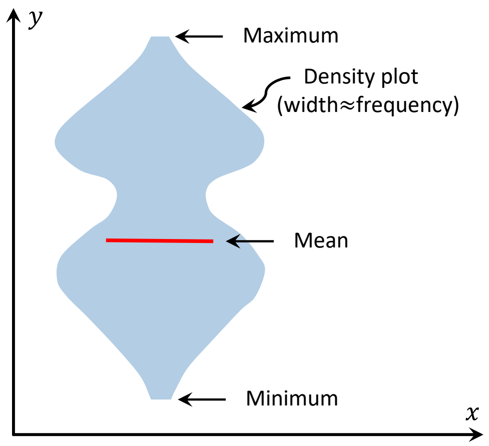


Figure S6: Interpretation of the violin plot

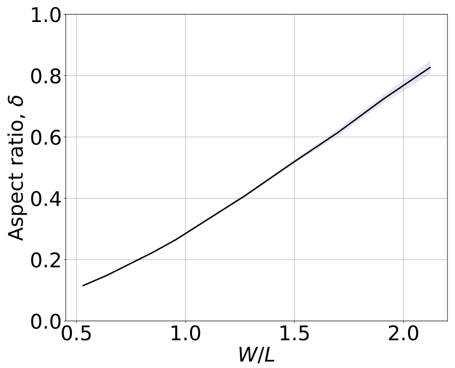


Figure S7: Cell aspect ratio versus the dimensionless channel width

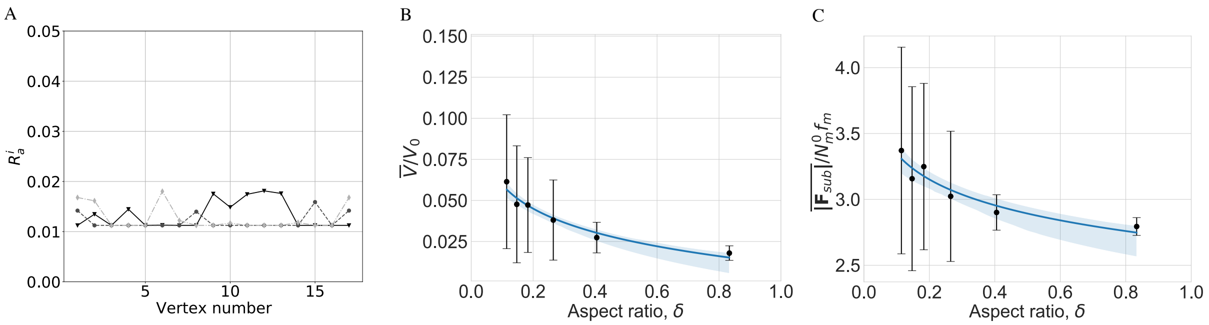


Figure S8: Effect of confinement on cell migration in the absence of persistent polarity. (A). Random distributions of the membrane-bound Rac1 in the active form across all vertices. Other forms of proteins are uniformly distributed. (B) $\overline{V}/V_0$ versus cell aspect ratios. The full line represents a linear regression model fit, along with a translucent 95% confidence interval band. Each data point and error bar represent the mean speed and standard deviation, respectively, calculated from more than 10 simulations. (C) The dimensionless mean net traction force ($\overline{|\mathbf{F}_{sub}|}/N_m^0 f_m$) as a function of cell aspect ratios.

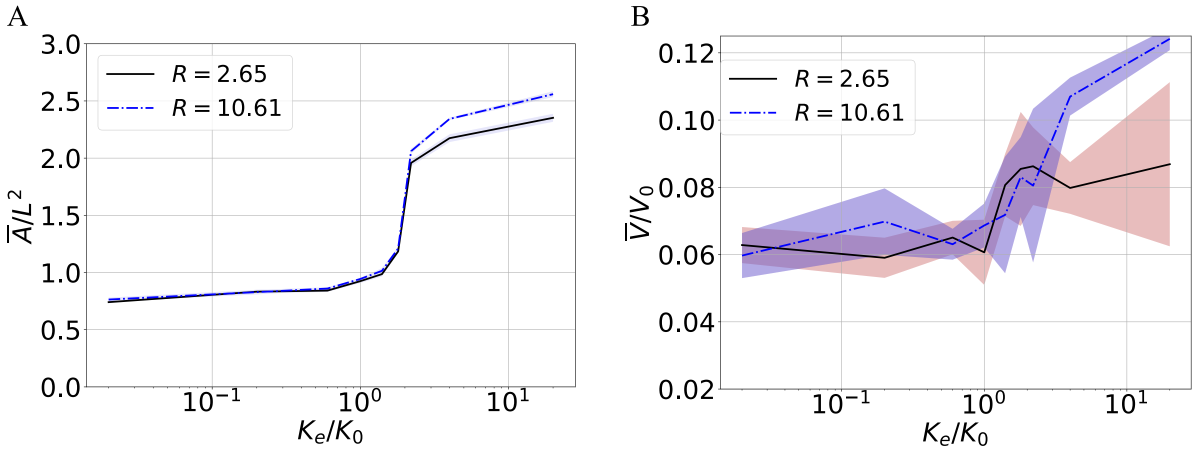


Figure S9: Influence of channel curvature decreases with smaller coefficient $\mu = 10^3\ /\mu m^2$. (A) The dimensionless mean spreading area $\overline{A}/L^2$ versus the dimensionless substrate elastic stiffness, $K_e/K_0$ (B) The dimensionless mean migration speed $\overline{V}/V_0$ and its standard deviation as a function of $K_e/K_0$ . Cells in (A) and (B) are confined in channels with the same channel width $W = 12\ \mu m$ but different radii ($R/L = 2.65$ and $R/L = 10.61$).

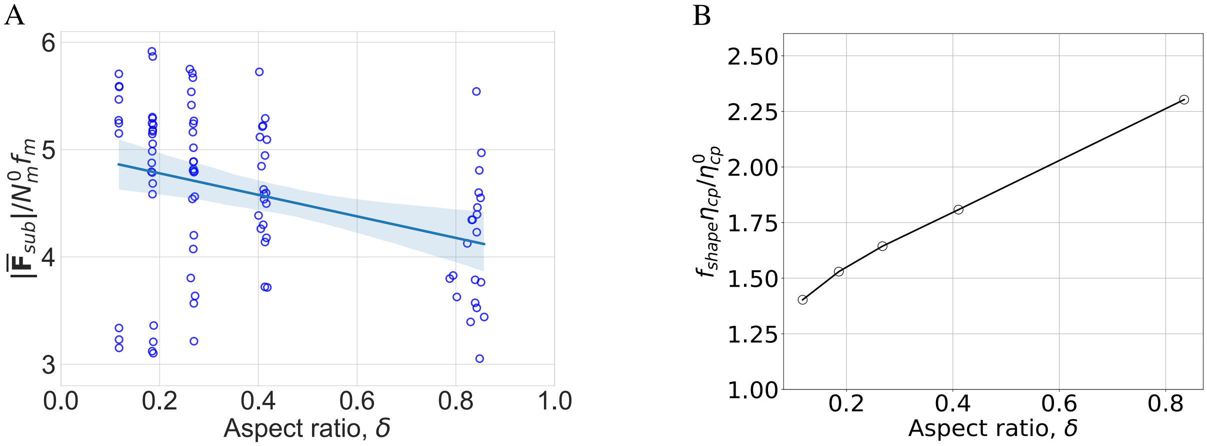


Figure S10: Changes of net traction forces and viscous drag on nucleus in less curved channels with different confinements. (A). The dimensionless net traction force $\overline{|\mathbf{F}_{sub}|}/N_m^0 f_m$, as a function of the cell aspect ratio for cells in channels with small curvature ($R/L = 10.65$). (B). Increase in nucleus viscous drag due to cell spreading, represented by $f_{shape}\eta_{cp}/\eta_{cp}^0$, versus the cell aspect ratio.

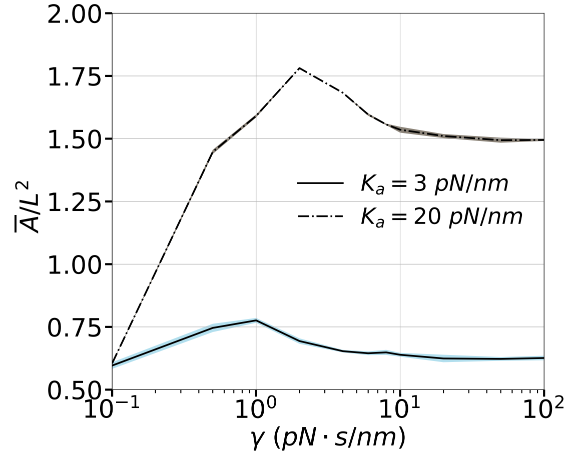


Figure S11: The dimensionless spreading area $\overline{A}/L^2$ and its standard deviation as a function of $\gamma$ for different additional stiffness $K_a$ ($n = 5$ .)

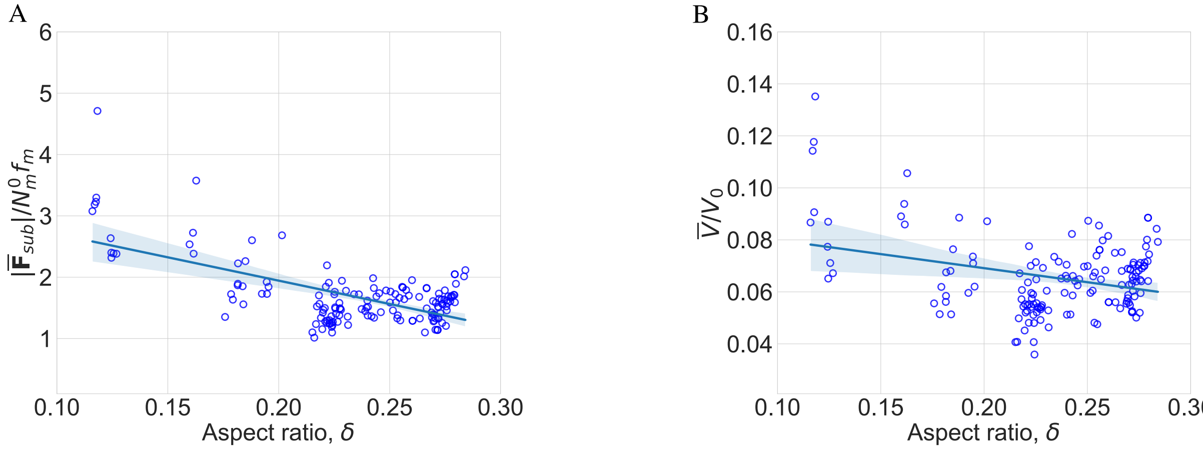


Figure S12: Relationship between aspect ratio, net traction force and mean dimensionless speed on viscoelastic substrates with strong confinement. (A) The dimensionless net traction force, $\overline{|\mathbf{F}_{sub}|}/N_m^0 f_m$, as a function of the cell aspect ratio. The full line represents a linear fit, along with a translucent 95% confidence interval band. Each dot represents a result obtained from a single simulation in heatmap Fig. 5B. (B) The dimensionless speed, $\overline{V}/V_0$, versus the cell aspect ratio.

Table S1: Model parameters used in the simulations.

| Symbol | Parameters | Dimensional | Dimensionless |
|---|---|---|---|
| $f_m$ | Single myosin motor stall force | 2.0 $pN$ [7, 8] | $1/100$ |
| $K_c$ | Clutch stiffness | 2.0 $pN/nm$ [8, 43] | $60\pi$ |
| $k_{on}^i$ | Rate constant of clutch association | 3.0 $s^{-1}$ [7] | $150\pi$ |
| $k_{r0}$ | Clutch unloaded off-rate | 0.25 $s^{-1}$ [7, 43] | $12.5\pi$ |
| $k_{c0}$ | Clutch unloaded catch-rate | 120 $s^{-1}$ [7, 43] | $6000\pi$ |
| $f_{c0}$ | Characteristic catch force | 0.5 $pN$ [7] | $1/400$ |
| $f_{r0}$ | Characteristic rupture force | 1.0 $pN$ [7] | $1/200$ |
| $V_0$ | Unloaded retrograde flow velocity | 120.0 $nm/s$ [8] | 1.0 |
| $V_p^0$ | Characteristic polymerization rate | 120.0 $nm/s$ [5] | 1.0 |
| $r_0$ | Cell radius | 3.0 $\mu m$ [5] | $1/2\pi$ |
| $r_{nuc}$ | Nucleus radius | 2.0 $\mu m$ [18, 20] | $1/3\pi$ |
| $K_m$ | Membrane stiffness | 10.0 $pN/\mu m$ [44] | $0.3\pi$ |
| $\eta_m^i$ | Viscosity of the surrounding medium | 50.0 $Pa \cdot s$ [23] | $0.18\pi$ |
| $K_{cs}^b$ | Baseline cytoskeletal stiffness | 1.25 $pN/\mu m$ [15, 45] | $3\pi/80$ |
| $\Delta K_{cs}$ | Stiffness increment | 2.8 $pN/\mu m$ [1] | $21\pi/250$ |
| $\beta$ | Strain-stiffening coefficient | 0.00185 [1] | |
| $\chi$ | Shape correlation coefficient | 0.0024 [1] | |
| $\zeta$ | Adhesion reinforcement coefficient | 0.5 $/pN$ , [15]) | 10 |
| $f_{cr}$ | Threshold force | 2.5 $pN$ | $1/80$ |
| $N_c^0$ | Reference molecular clutch number | 100 [7] | 100 |
| $N_m^0$ | Reference myosin motor number | 100 [7] | 100 |
| $M_R^+$, $M_\rho^+$ | Rac1, RhoA membrane association rate | 0.02 $s^{-1}$ [46] | $\pi$ |
| $M_R^-$, $M_\rho^-$ | Rac1, RhoA membrane dissociation rate | 0.02 $s^{-1}$ [46] | $\pi$ |
| $K_b^+$, $\kappa_b^+$ | Baseline Rac1, RhoA activation rates | 0.3 $s^{-1}$ | $15\pi$ |
| $K^-$ | Rac1 deactivation rate ($I_R^i = K^-$) | 0.8 $s^{-1}$ [34, 37] | $40\pi$ |
| $\kappa^-$ | RhoA deactivation rate ($I_\rho^i = \kappa^-$) | 0.8 $s^{-1}$ [34, 37] | $40\pi$ |
| $R_0$, $\rho_0$ | Reference level of the active Rac1 and RhoA | 3/160 | 3/160 |
| $\theta_0$ | Vertex angle at $t = 0$ | $7\pi/8$ | $7\pi/8$ |
| $c_0$ | Ratio of the threshold angle | 0.3 | 0.3 |
| $\alpha_R$, $\alpha_\rho$ | Positive feedback rate on Rac1 and RhoA | 0.3 $s^{-1}$ [37] | $15\pi$ |
| $\beta_R$ | Rate of Rac1 inhibition by RhoA | 0.3 $s^{-1}$ [47] | $15\pi$ |
| $\beta_\rho$ | Rate of RhoA inhibition by Rac1 | 0.3 $s^{-1}$ [47] | $15\pi$ |
| $D$ | Diffusivity on the membrane | 0.01 $\mu m^2/s$ [37] | $1/72\pi$ |
| $\eta_{cp}^0$ | Reference cytoplasm viscosity | 860 $Pa \cdot s$ [32] | $3.096\pi$ |
| $\Delta\eta_{cp}$ | Cytoplasm viscosity increment | 36 $Pa \cdot s$ [32] | $81\pi/625$ |
| $\epsilon$ | Coefficient in confined channels | 0.5 (Fig.S1) | 0.5 |
| $\mu$ | Coefficient in curved channels | $4 \times 10^4$ $\mu m^2$ (Fig.S3) | $10000/9\pi$ |
| $\phi_0$ | Threshold angle between cytoskeleton elements | $\pi/30$ | |
| $K_{rp}$ | Penalty parameter | $10^3$ | 5 |
